# Invasion and maintenance of meiotic drivers in populations of ascomycete fungi

**DOI:** 10.1101/2020.04.06.026989

**Authors:** Ivain Martinossi-Allibert, Carl Veller, S. Lorena Ament-Velásquez, Aaron A. Vogan, Claus Rueffler, Hanna Johannesson

## Abstract

Meiotic drivers are selfish genetic elements that are able to become over-represented among the products of meiosis. This transmission advantage makes it possible for them to spread in a population even when they impose fitness costs on their host organisms. Whether a meiotic driver can invade a population, and subsequently reach fixation or coexist in a stable polymorphism, depends on the one hand on the biology of the host organism, including its life-cycle, mating system, and population structure, and on the other hand on the specific fitness effects of the driving allele on the host. Here, we present a population genetic model for spore killing, a type of drive specific to fungi. We show how ploidy level, rate of selfing, and efficiency of spore killing affect the invasion probability of a driving allele and the conditions for its stable coexistence with the non-driving allele. Our model can be adapted to different fungal life-cycles, and is applied here to two well-studied genera of filamentous ascomycetes known to harbor spore killing elements, *Neurospora* and *Podospora*. We discuss our results in the light of recent empirical findings for these two systems.

## 1 Introduction

It is often assumed in genetics that the two copies of a gene in a diploid genome are represented equally among the products of meiosis — this is Mendel’s first law (Lyttle, 1993). However, some genetic elements called meiotic drivers (MDs) are able to distort meiosis and become over-represented among meiotic products (Sandler and Novitski, 1957; Burt and Trivers, 2006). Due to their ability to distort meiosis, MDs gain a selective advantage that allows them to increase in frequency in a population even when they impose fitness costs on their host organism (Hamilton, 1967; Akbari et al., 2013; Pinzone and Dyer, 2013; Kyrou et al., 2018). The ensuing genetic conflict between MDs and their hosts is known to affect several evolutionary processes (Rice, 2013). For example, rapid co-evolution between MDs and counteracting genes, called suppressors, can accelerate speciation by creating genetic incompatibilities between recently separated populations (Frank, 1991), as well as shape genetic architecture in other important ways (Henikoff et al., 2001; Hurst and Werren, 2001; Werren, 2011).

MDs were discovered as early as 1928 (Sandler and Novitski, 1957) and have been studied extensively since then. Early empirical observations were closely followed by theoretical work aimed at understanding the unique population genetic behavior of these selfish genetic elements (see for example Hiraizumi, 1962; Lewontin and Dunn, 1960; Lewontin, 1968, on the *t-haplotype* in mice). Theoretical work has focused on two key aspects of meiotic drive dynamics: under what conditions can a MD (i) invade a population and (ii) coexist at a stable equilibrium with a non-driving allele? These questions have been investigated with reference to a wide variety of species harboring MDs (e.g. Lewontin and Dunn, 1960; Fishman and Kelly, 2015; Brand et al., 2015; Hall and Dawe, 2018), which has revealed some general patterns of MD dynamics. First, since MDs are over-represented among successful meiotic products, theory predicts that, in the absence of counteracting forces, they should increase in frequency and reach fixation. However, the presence of suppressor alleles or fitness costs associated with the MD can bring the invasion process to a halt, leading ultimately to either the loss of the MD or prolonged coexistence with a non-driving allele. A typical condition for coexistence appears to be the presence of strong recessive fitness costs to the driver (or the haplotype on which it lies), allowing invasion of the MD but not fixation (e.g. Fishman and Kelly, 2015; Lewontin and Dunn, 1960; Holman et al., 2015). These general principles, however, are far from encompassing the complexity and diversity of MD dynamics. Indeed, although all MDs share the feature of distorting Mendelian segregation, the diversity of their modes of action as well as differences in the life cycles of their hosts make insights from one system often not applicable to others.

Here, we focus on a type of MD that has been relatively neglected by theoreticians (but see Nauta and Hoekstra, 1993): spore killers MDs in ascomycete fungi. Spore killers are found in several species of ascomycete fungi (Padieu and Bernet, 1967; Turner and Perkins, 1979; Zanders et al., 2014) and there are good reasons to expect that their dynamics differ from that of well-known meiotic drivers in plant and animal systems. The diversity of life cycles found among ascomycetes, together with the distinct mechanism of drive of spore killers (Núñez et al., 2018), could influence the dynamics of these MDs in yet unknown ways.

Meiotic drive in animals and plants occurs during gametogenesis, and can be categorized into either female drive or male drive (reviewed in Burt and Trivers, 2006; Lindholm et al., 2016). In female drive, as observed, for example, in maize (Buckler et al., 1999), monkeyflowers (Fishman and Saunders, 2008), and mice (Didion et al., 2015), the MD takes advantage of the asymmetry of female meiosis by preferentially segregating to the functional egg cell (or macrospore). In contrast, in male drive, the MD kills or disables gametes that do not carry it. Well-known examples of male drive are the *t-haplotype* in *Mus musculus* (Silver, 1985) and *SD* in *Drosophila melanogaster* (Larracuente and Presgraves, 2012). Spore killing may appear, at first, to be similar to male drive because it also entails the killing of meiotic products that do not carry the MD (Raju, 1994). However, because the dominant phase of the fungal life cycle is haploid, meiosis does not correspond to gametogenesis but instead to the production of offspring (haploid spores). Spore killing occurs when the spores produced by a single meiosis are packaged together in a sac (‘ascus’) prior to their dispersal. Their proximity in the ascus allows spores carrying the MD to disable spores that do not carry it. Another specificity of spore killing is that it can affect all individuals in the population, instead of being limited to one sex.

The nature of the selective advantage of a MD depends on its mechanism of drive. These fall into two categories: absolute drive and relative drive (Lyttle, 1991). In relative drive the MD only increases in relative frequency by reducing the number of gametes not carrying the MD while absolute drivers increase the number of gametes carrying the MD in absolute numbers (Figure 1). The selective advantage of a purely relative driver is small during the early stages of invasion when the driver is at low frequency (as pointed out by Nauta and Hoekstra, 1993), while absolute drivers have a high initial selective advantage because they increase in absolute copy number when driving. This distinction can have a great impact on the likelihood of invasion of a MD, and it is therefore important to understand to which category spore killers can belong. Female drive can be purely absolute drive because the MD can be more likely to end up in the functional egg (or macrospore) than the alternative allele, with no reduction in the total number of eggs produced. In this way, the MD can increase in absolute number of copies (Figure 1) with little or no fitness costs to the host. In contrast, male drive can be purely relative drive if its action results exclusively in the killing of gametes carrying the non-driving alleles: the MD increases in frequency relative to the non-driver allele, but does not increase in absolute copy number (Figure 1).

**Figure 1:**
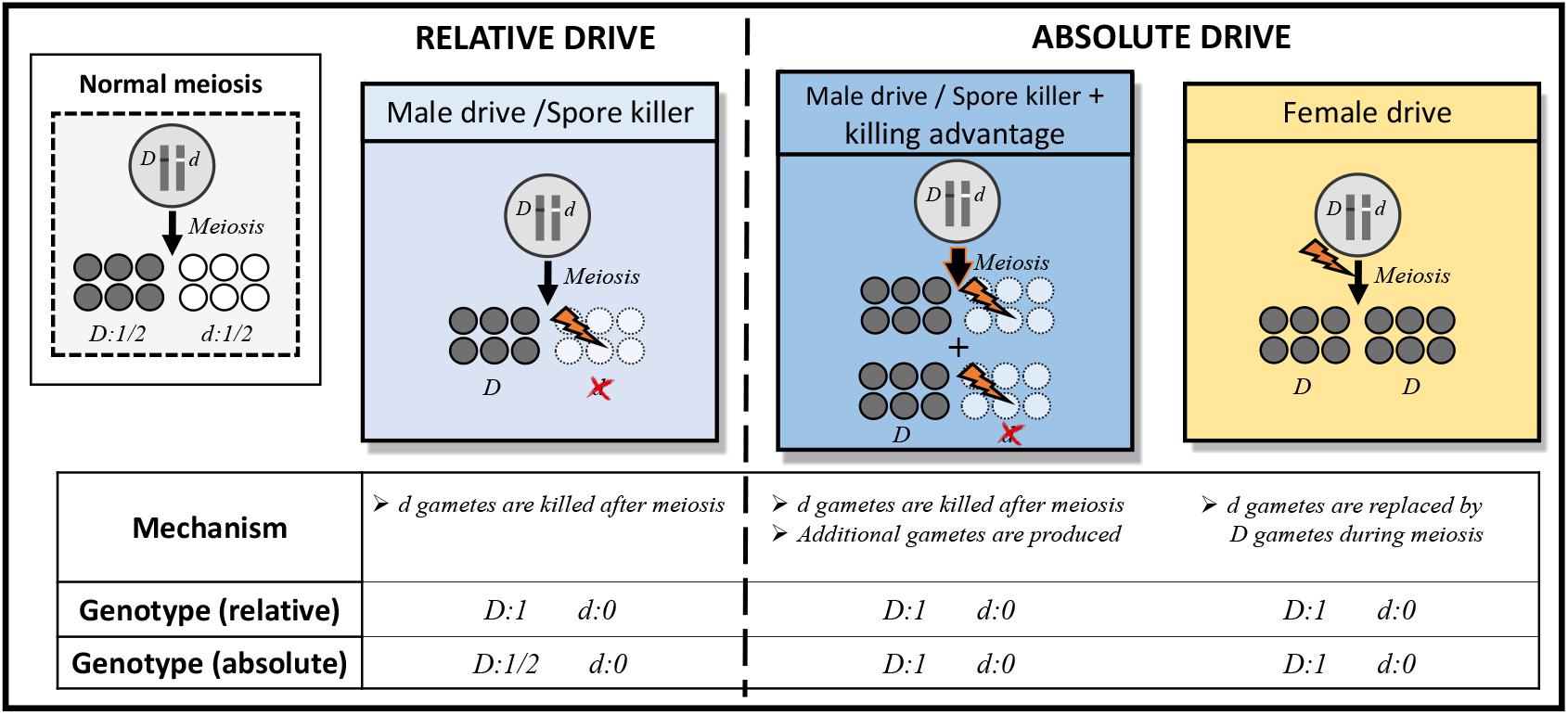
Relative and absolute meiotic drive. The organism is diploid, and heterozygous for alleles *D* (driving allele) and *d* (sensitive allele) at a given locus. The figure shows how relative and absolute drive differ from normal meiosis.

Male drive can enjoy an absolute advantage if the killing of sensitive meiotic products results in either the production of additional meiotic products or higher absolute fitness of surviving ones. We refer to such benefits generally as *killing advantage* (Figure 1). In male drive, killing advantage (also called ‘compensation’) can arise when fertility does not scale linearly with the number of sperm produced (Hartl, 1970): halving the sperm count may not half the offspring number, so that a MD that kills all sperm that carry the sensitive allele can end up in a greater number of offspring than expected for a neutral allele in a heterozygous parent. Killing advantage in male drive can also evolve as an adaptive response of the host organism, for example, heterozygous carriers of the MD could produce additional gametes to compensate for those killed by the MD (e.g. in the stalk-eyed fly *Teleopsis dalamanni* ; Meade et al., 2019).

At a first glance, spore killing in fungi appears to be a purely relative drive because spore killers directly eliminate some proportion of their host’s offspring. However, killing advantage may be possible in spore killers as well, with mechanisms somewhat similar to certain forms of killing advantage in male drive. For instance, the killing of sibling spores may alleviate local competition for resources, thus increasing the viability of surviving spores. Alternatively, the parent might have a lower resource expenditure on killed spores, and thus be able to produce additional spores to compensate for the loss (Nauta and Hoekstra, 1993; Lindholm et al., 2016). In both scenarios, the killing advantage would provide the spore killer with a fitness benefit that makes it more akin to an absolute driver. In natural systems, it is not known whether a killing advantage exists for spore killers—and of what type—except for the case of *Podospora anserina*, where the production of additional spores in response to spore killing has been documented (Vogan et al., 2020).

In this study, we develop a population genetic model that allows us to address the two central questions of theoretical meiotic drive studies in the relatively uncharted realm of spore killers: Under what conditions can a spore killing MD (i) invade and (ii) coexist with a sensitive allele? We build a single-locus model of spore killing. In considering a single-locus model, we ignore—in agreement with recent empirical findings (Nuckolls et al., 2017; Hu et al., 2017; Vogan et al., 2019; Svedberg et al., 2020)—the possibility of recombination between the killing and antidote functions of the spore killer, which was addressed in the only other theoretical work we know of on spore killers (Nauta and Hoekstra, 1993). We cover the diversity of ascomycete life cycles by adapting our model to heterothallic *Neurospora* species, which are representative of typical ascomycete fungi, and *Podospora anserina*, which has evolved a more complex life cycle (Raju and Perkins, 1994). In our study, we investigate the influence of fitness costs associated with the MD, killing efficiency, and killing advantage on the dynamics of the spore killer. For *P. anserina*, we also consider the impact of two additional parameters that are of interest in this case: the selfing rate and the recombination rate between the spore killer locus and the centromere. We investigate these questions by performing a stability analysis of the equilibria of the recursion equation describing the dynamics of the spore killer allele and with stochastic simulations to assess the impact of drift on the invasion probability of a spore killer.

## 2 The model

We study allelic dynamics at a single locus in a diploid organism with non-overlapping generations. Two alleles segregate at the focal locus: the spore killer allele *D* and the sensitive non-killing allele *d*. The organismal life cycles that we consider correspond to those of filamentous ascomycetes of the genera *Neurospora* and *Podospora*. Both taxa are model systems in fungal genetics and are known to harbor spore killing elements (e.g. Silar, 2013; Vogan et al., 2019; Svedberg et al., 2020). Moreover, the life cycle of heterothallic *Neurospora* species is representative of a typical filamentous ascomycetes and the life cycle of *P. anserina* represents pseudohomothallism, an alternative reproductive strategy that has evolved many times across ascomycetes with a variety of genetic mechanisms (Raju, 1994). These two species can therefore capture a wide range of the variety of ascomycete life cycles. In our analysis, we first assume that the population is sufficiently large that drift can be ignored. Under this assumption, we determine the parameter combinations that permit invasion of the spore killer allele *D*, and then ask under what further conditions invasion of *D* results in its fixation or its stable coexistence with *d*. We then relax the assumption of an infinitely large population to explore the role of drift in the early phase of invasion of *D*.

### 2.1 The *Neurospora* model

#### 2.1.1 *Neurospora* life cycle and model parameters

In this section, we focus on the life cycle of heterothallic *Neurospora* species, such as *N. sitophila* and *N. crassa*, where different mating types occur in different individuals. Since individuals in these species are sexually self-incompatible, we assume random mating and therefore Hardy-Weinberg proportions, although we acknowledge that inbreeding by sib-mating is a possibility in *Neurospora*. The simple structure of the life cycle is summarized in Figure 2a. A haploid vegetative stage leads to the production of haploid gametes and after fusion of gametes of opposite mating types a short-lived diploid zygote is formed that quickly undergoes meiosis. Meiosis followed by a post-meiotic mitosis results in the formation of an eight-spore ascus. Spores disperse and grow to become the vegetative stage of the next generation. When an ascus originates from a diploid individual that is heterozygous at the spore killer locus, spore killing occurs and spores of genotype *d* are killed with probability *e*, so that a proportion 1 − *e* of such spores survive. We refer to *e* as the killing efficiency. When a killing event occurs, all surviving spores within the ascus (all *D*-spores and an expected fraction 1 − *e* of *d*-spores) benefit from a killing advantage *a* < 1. At each stage of the life cycle, carrying the *D* allele is associated with a viability cost represented by parameters *z* for the diploid stage and *g* for the haploid stage. *DD* diploids suffer a cost *z*, while *Dd* heterozygotes suffer a cost *zh*_*z*_, where *h*_*z*_ is the dominance coefficient. Figure 2a shows at what stage the different parameters apply in the life cycle of *Neurospora*. The assumption of random mating allows us to investigate the model by following a single variable, the frequency of the *D* allele among the pool of (randomly mating) gametes.

**Figure 2:**
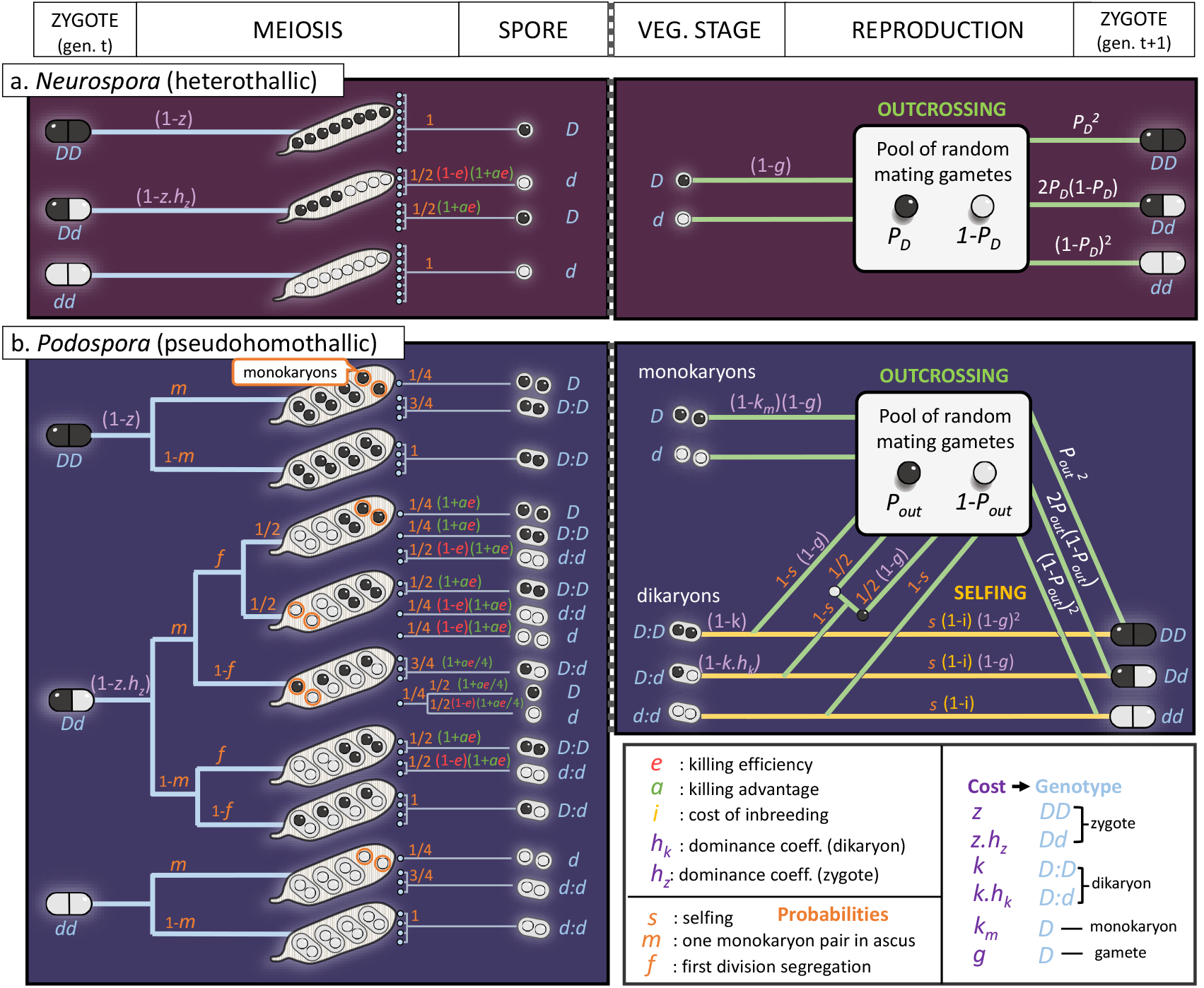
Life cycle diagrams for (a) heterothallic *Neurospora* species and (b) *P. anserina*. Both life cycles are presented from zygote to zygote. The left panel shows the meiosis stage occurring shortly after zygote formation and asci representing all possible segregation patterns of *D* and *d* from the three possible diploid genotypes. The right panel shows the reproductive stage with the outcrossing and selfing modes (the latter one only in *Podospora*) of reproduction, leading to fertilization and the formation of the diploid zygotes of the next generation. The recursion equations can be constructed by following the lines of the life cycle diagram and multiplying each genotype frequency with the costs and probabilities that apply to it as indicated along the lines. Specifically, orange symbols represent probabilities of alternative events (for example, selfing occurring with probability *s* versus outcrossing occurring with probability 1 − *s*). Purple symbols represent potential fitness costs associated with carrying the spore killer allele *D*, while the cost of inbreeding (*i*) at all loci, associated with the selfing branch of reproduction is represented in yellow. Furthermore, green symbols represent a killing advantage (*a*) associated with spore killing and killing efficiency (*e*) is colored in red.

#### 2.1.2 Recursion equation for *Neurospora*

Let *p*_*D*_ be the frequency of allele *D* in the gamete pool in the current generation, and 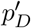 its frequency in the next generation. Then

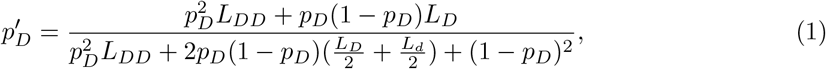

where

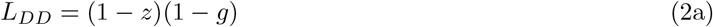

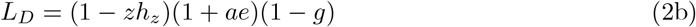

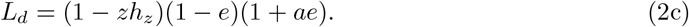

Here, *L*_*DD*_ represents the fitness of a *D* nucleus in a *DD* zygote, *L*_*D*_ the fitness of a *D* nucleus in a *Dd* zygote, and *L*_*d*_ the fitness of a *d* nucleus in a *Dd* zygote, which, in this last case, includes the costs of being killed during the spore killing event that follows meiosis.

### 2.2 The *Podospora* model

#### 2.2.1 *Podospora* life cycle and model parameters

Figure 2b shows a schematic representation of the more complex life cycle of *P. anserina*, from which we derive a set of recursion equations describing the change in frequency of the spore killer allele *D*. A simplified biological representation of the life cycle is shown in Figure S22. Additional complexities present in *P. anserina* include a mainly dikaryotic spore packaging and vegetative stage and alternative selfing and outcrossing reproductive strategies. The life cycle starts with a zygote (left panel of Figure 2b) that shortly after undergoes meiosis followed by one mitosis. This results in a single ascus containing eight haploid nuclei. These nuclei are then packaged into pairs, each pair enclosed within a dikaryotic spore (two haploid nuclei in the same cytoplasm), or remain isolated, resulting in monokaryotic spores (haploid). In *P. anserina*, the frequency of monokaryotic spores formed in natural strains is low (van der Gaag, 2005), so we assume that each ascus can contain either one pair of monokaryotic spores (with probability *m*) or none (with probability 1− *m*). Heterozygous diploid cells *Dd* can result in the formation of either heteroallelic *Dd* or homoallelic *DD* and *dd* dikaryotic spores, depending on whether the drive locus segregates reductionally (first division segregation, or FDS) or equationally (second division segregation, or SDS) at the first meiotic division (Rizet and Engelmann, 1949). FDS at the drive locus occurs when there is no crossover, or alternatively an even number of crossovers, between the drive locus and the centromere. SDS occurs otherwise (see meiosis diagram in Figure S22). We assume that FDS occurs at the spore killer locus with probability *f* and results in two homoallelic spores of each genotype, while SDS occurs with probability 1 − *f* and results in four heteroallelic spores (see Figure 2b). When monokaryotic spores of genotype *d* or dikaryotic spores of genotype *dd* share an ascus with spores of genotype *D, DD* or *Dd*, they are killed with efficiency *e*. Dikaryotic spores of the *Dd* genotype are not affected by spore killing, because the *D* nucleus protects the entire spore from being killed (Grognet et al., 2014; Padieu and Bernet, 1967). The frequencies of the different types of spores after meiosis are denoted by *M*_*DD*_, *M*_*Dd*_, *M*_*dd*_, *M*_*D*_ and *M*_*d*_, respectively.

After meiosis, both dikaryotic and monokaryotic spores germinate and form a mycelium, which is the vegetative stage of the life cycle. We assume that dikaryons and monokaryons have the same growth rate during this stage. The vegetative stage is followed by the reproductive stage, which is represented in the right panel of Figure 2. Gametes produced by both dikaryons and monokaryons enter a common pool where random mating results in diploid zygotes. In addition, dikaryons can self, which they do with probability *s*. Selfing in *P. anserina* can occur only when a dikaryon carries nuclei of the two different mating types. This is always the case because the mating-type locus undergoes SDS during meiosis, making dikaryons automatically heteroallelic for the mating-type locus (in nature, the probability of SDS of the mating-type locus is very close to 100% in *P. anserina*, Rizet and Engelmann, 1949). As a consequence of this mating type constraint, heteroallelic dikaryons of genotype *Dd* can only produce heterozygous diploid zygotes *Dd* through selfing, because each of the *D* and *d* alleles is associated with a particular mating type, thus preventing the formation of *DD* or *dd* individuals. We note that, although this particular type of selfing maintains heterozygous genotypes during reproduction, it still results in a loss of heterozygosity because heterozygous genotypes can be broken apart by FDS during meiosis. At each stage of the life cycle, our model includes the possibility of fitness costs resulting in reduced viability associated with the spore-killing allele *D* (see Figure 2). Costs are the same as for the *Neurospora* model, with the addition of a cost *k* in the dikaryotic stage and its associated dominance coefficient *h*_*k*_.

#### 2.2.2 Recursion equation for a simplified *Podospora* model

We first present a simplified *Podospora* model, which builds on the *Neurospora* model with the addition of a dikaryotic cost parameter *k* and a rate of FDS *f*. This simplified model does not yet include selfing or monokaryons, and therefore does not fully represent the biology of *Podospora*, but it is useful in allowing us to derive simple analytical results that help clarify the interpretation of the full *Podospora* model introduced in the next section. Thus, assuming that the rate of selfing is zero (*s* = 0) and that monokaryons do not occur (*m* = 0), recursion (1) from the *Neurospora* model applies to this simplified version of the *Podospora* model with the following modifications:

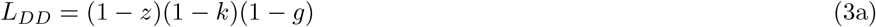

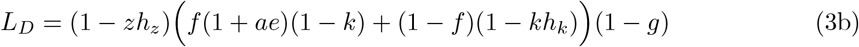

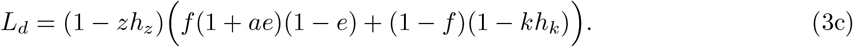

#### 2.2.3 Recursion equations for the complete *Podospora model*

In the full *Podospora* model, the occurrence of selfing implies that mating is not random, and so we need to track the frequencies of the diploid genotypes *DD, Dd* and *dd*. Each generation, some individuals from each genotype are produced through selfing and some through outcrossing. Thus, the change in frequency for the three genotypes *DD, Dd* and *dd* is given by

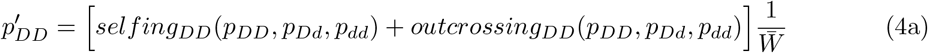

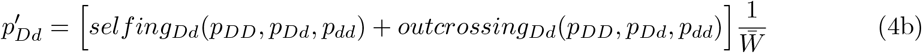

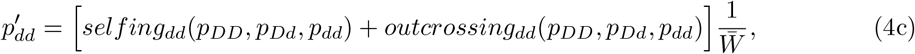

where the frequencies of a given genotype in the current and next generation are indicated by *p* and *p*′ respectively, with genotypes indicated by subscripts, and 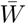 representing mean fitness.

The treatment of selfing and outcrossing is inspired by a plant population genetics model with self fertilization (Holsinger et al., 1984). Beginning with the selfing part of the life cycle (right-hand panel of Figure 2), we start from the frequencies of the three possible dikaryotic genotypes after meiosis, *M*_*DD*_, *M*_*Dd*_ and *M*_*dd*_. Dikaryons may pay fitness costs if they carry one or two copies of the spore killer (respectively *kh*_*k*_ or *k*). Genotype frequencies are then adjusted by the selfing probability *s*, and all genotypes are exposed to a selfing cost *i* due to inbreeding. Finally, because gametes are also produced during the selfing process, individuals carrying the spore killer genotype are exposed to gametic costs *g* during selfing (as well as during outcrossing), resulting in the survival probabilities (1 − *g*)^2^ and 1 − *g* for *DD* and *Dd* dikaryons, respectively. We can now write the selfing contribution to next generation’s diploid genotype as

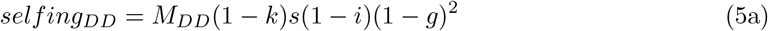

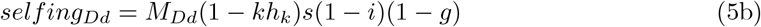

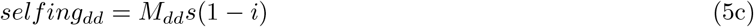

For the outcrossing part of the life cycle, random mating is assumed. We denote by *p*_out_ the frequency of the spore-killing allele *D* in the pool of randomly mating gametes. It is important to note that *p*_out_ only represents a frequency within the outcrossing fraction of the total population, denoted by *T*_out_. More precisely, *T*_out_ consists the fraction of gametes that engage in outcrossing, which are all gametes from monokaryotic individuals (potentially reduced due to fitness costs *k*_*m*_ for monokaryons carrying the *D* allele) together with the fraction 1 − *s* of outcrossing gametes from dikaryons, all discounted by the appropriate reduction in viability due to costs. Genotypes produced by outcrossing follow Hardy-Weinberg proportions and are weighted by *T*_out_ to represent valid frequencies in the total population,

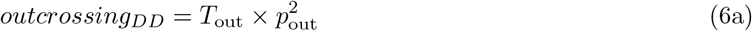

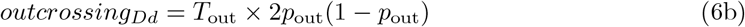

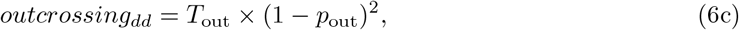

where

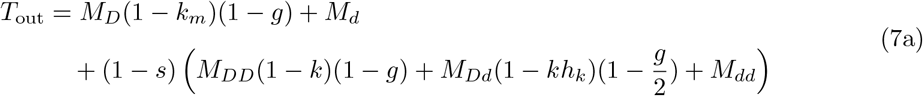

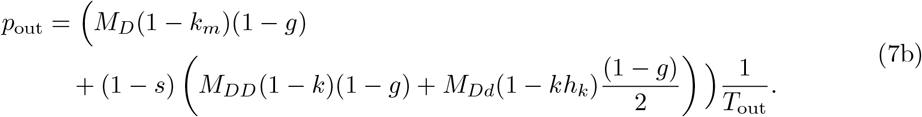

The expressions for the genotype frequencies after meiosis are given by

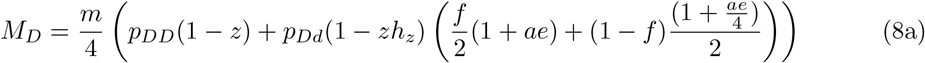

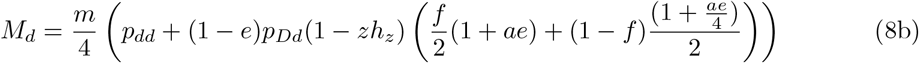

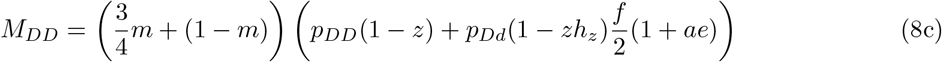

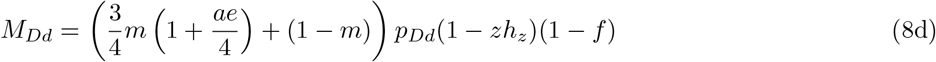

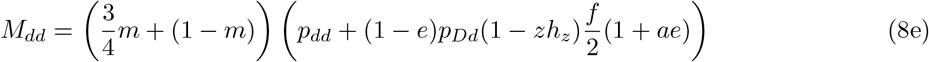

and can be derived from the left-hand panel of Figure 2. Fitness costs can affect diploid zygotes, reducing the initial frequencies by the factors 1 − *z* and 1 − *zh*_*z*_ for *DD* and *Dd* genotypes, respectively. When a pair of monokaryons are formed (with probability *m*), they represent 1*/*4 of the nuclei in their ascus. Consequently, monokaryotic spores represent a fraction *m/*4 of the initial diploid frequencies, and dikaryons represent a fraction 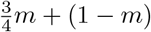. Spore killing affects monokaryotic spores of the *d* genotype and dikaryotic spores of the *dd* genotype originating from *Dd* diploids. In these cases, killing occurs with efficiency *e*. Thus, a proportion 1− *e* of the sensitive spores exposed to killing survive. Importantly, monokaryotic *d*-spores are affected under both first-and second-division segregation, while dikaryotic spores are affected only when homoallelic (*dd*), and therefore only under first-division segregation (which occurs with probability *f*). Only when the killing efficiency is maximal (*e* = 1) are all sensitive spores that are exposed to the *D* allele killed. In asci in which spore killing occurs, all surviving spores can benefit from a killing advantage regardless of their genotype. The killing advantage is likely to originate from additional resources made available due to spores being killed and is therefore assumed to be proportional to the number of killed spores. For this reason, the killing advantage is weighted by the killing efficiency, providing a benefit 1 + *ae* to surviving spores. The killing advantage benefiting monokaryotic spores of an ascus in which the killer locus underwent SDS is special. In this case, only one nucleus per ascus can be killed in contrast to the four nuclei that are killed under FDS. When this occurs, the killing advantage is 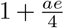. Finally, 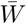 is the sum of the numerators on the right-hand side of equation (4).

#### 2.2.4 Expected parameter range for *Podospora*

For this system, it is unknown whether spore-killing alleles impose fitness costs on the carrier and if, and to what extent, a killing advantage exists. Therefore, we study the broadest possible range of fitness costs for each stage of the life cycle, from 0 (no costs) to 1 (fully lethal), and a wide range of killing advantages from 0 (no benefit) to 1 (equivalent to all killed spores being replaced). Also the rate of selfing in natural populations of *Podospora* is not known. However, the propensity of *Podospora* species to self in laboratory conditions together with low overall levels of genetic diversity (Vogan et al., 2019) indicate that selfing may be frequent. Therefore, we study the effect of selfing rates ranging from 0 to 95%. Known spore killers in *Podospora* undergo FDS in 30-100% of meioses, depending on the variants (van der Gaag et al., 2000; Vogan et al., 2019). We cover this range by studying the probabilities 0.25, 0.5 and 1 of FDS. Under natural conditions, the occurrence of asci containing monokaryons can vary between 0% and 6% (Esser, 1974; van der Gaag, 2005). We therefore analyse the model without monokaryons first, and then with monokaryons occurring in 5% of asci to cover the natural range, and finally in 50% of asci to make the effect of monokaryons on spore killing dynamics more visible. Finally, the killing efficiency *e* is believed to be high in *Podospora* (Vogan et al., 2019). Hence, we focus on the case with *e* = 1, and only briefly explore incomplete killing (*e* < 1) to contrast its effect with that of the probability of FDS *f* < 1.

### 2.3 Methods

We first analyse the deterministic recursions to characterise the parameter combinations that permit invasion with subsequent fixation of *D*, or invasion with subsequent stable polymorphism. To this end, we identify the equilibria of the recursions and determine their stability. Stability is determined based on a linear stability analysis of the one-dimensional system in the cases of *Neurospora* and simplified *Podospora* without selfing (Otto and Day, 2011, pp. 163-172) or the two-dimensional system in the case of *P. anserina* with selfing (Otto and Day, 2011, pp. 316-320). For the one-dimensional models we obtain analytical results that allow us to identify exact conditions for invasion and stable polymorphism as functions of the model parameters. In the case of the complete *P. anserina* model, we can only solve for equilibria and stability for given parameter values, and we therefore conduct parameter sweeps.

In order to determine the invasion probability of a spore-killing allele in a finite population, we analyse a stochastic version of the model accounting for drift. In this analysis, the recursions define the sampling probabilities of a Wright-Fisher process, with sampling occurring at the stage of zygote formation. For each set of parameter values, the invasion probability is estimated as the proportion of 1000 simulation runs in which the spore killer achieves the equilibrium frequency of the deterministic system (fixation or stable polymorphism). A stochastic simulation is considered to have reached an internal equilibrium if allele frequencies fluctuate around the same value for at least 1000 generations. The invasion probability is taken to be zero whenever the deterministic model does not allow for invasion.

## 3 Results

### 3.1 Neurospora

#### 3.1.1 Deterministic model

The dynamics of the spore-killing allele *D* in *Neurospora*, as described by equation (1), can be analyzed analytically. Solving this equation for its equilibria, we obtain 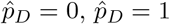, and

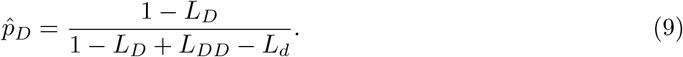

This last equilibrium is valid (i.e., 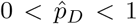) if *L*_*D*_ < 1 and *L*_*DD*_ > *L*_*d*_, or if *L*_*D*_ > 1 and *L*_*DD*_ < *L*_*d*_. Below we discuss how the conditions for invasion, fixation and coexistence of the killer allele *D* are affected by its killing efficiency, the killing advantage, and fitness costs and their degree of dominance. Figure 3 summarizes these results for the case of diploid fitness costs *z* while the results for the case of haploid costs *g* are shown in Figure S1.

**Figure 3:**
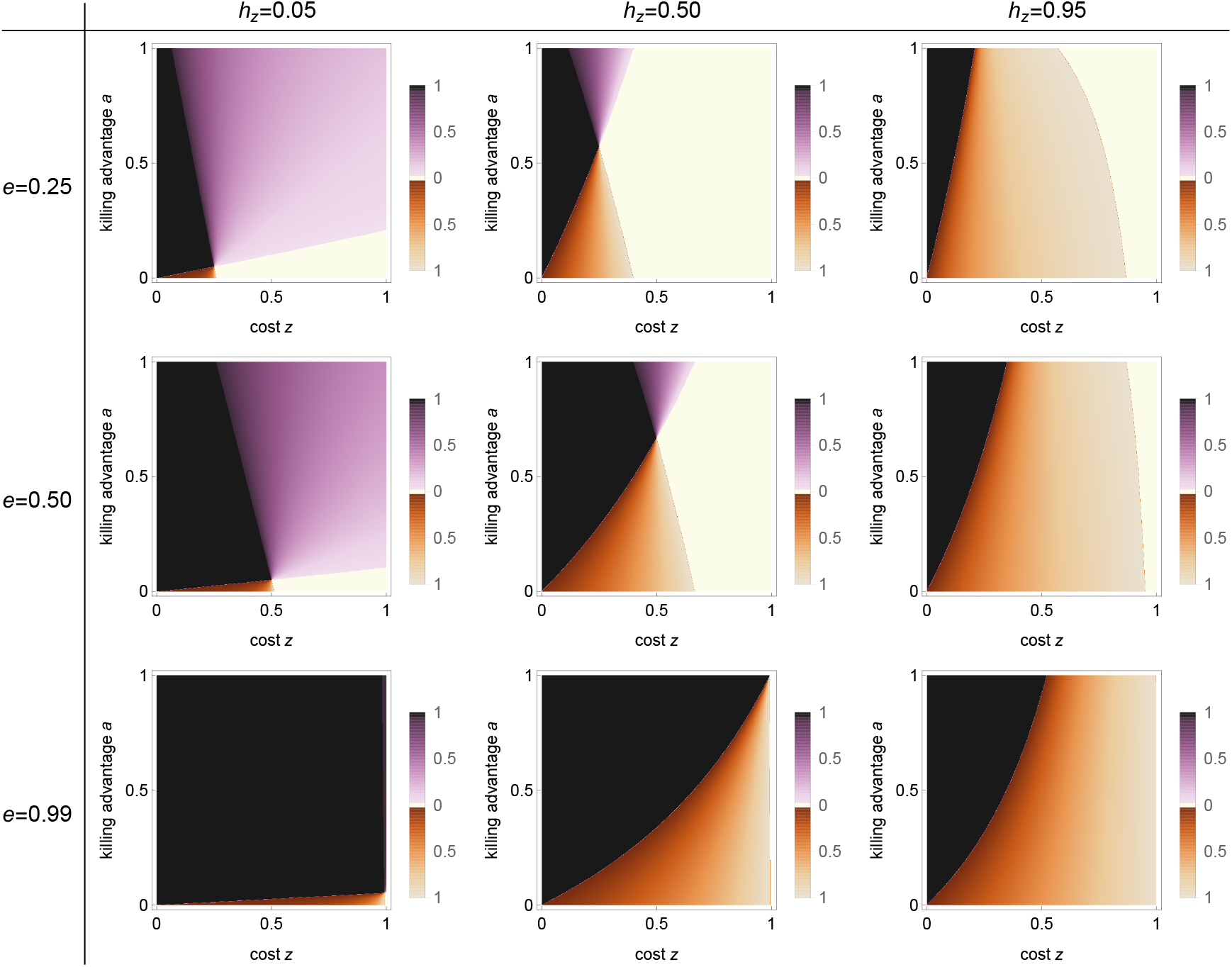
Bifurcation analysis of the *Neurospora* model with diploid fitness costs *z*. The bifurcation parameters are diploid fitness costs *z* and their dominance *h*_*z*_, killing advantage *a*, and killing efficiency *e*. Extinction 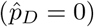 and fixation 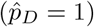 of the killer allele *D* always constitute equilibria. Additionally, one interior equilibrium is possible. Parameter regions are color-coded as follows: **white**, *D* cannot invade and 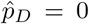 is a globally stable equilibrium; **black**, *D* can invade and reach fixation, 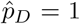 is a globally stable equilibrium; **purple**, *D* can invade but cannot reach fixation, instead coexisting with the non-killing allele *d* at a globally stable interior equilibrium 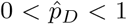, whose value is given by the shade of purple; **brown**, the two boundary equilibria 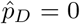 and 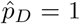 are stable, with their basins of attraction separated by an unstable interior equilibrium 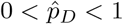 whose value is given by the shade of brown. Each panel is based on 1000×1000 parameter combinations. Gametic fitness costs *g* = 0.

A linear stability analysis shows that the killing allele *D* can invade (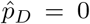 unstable) if *L*_*D*_ > 1. From equation (2b) we can see that this condition is met when

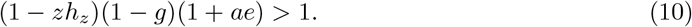

Thus, invasion is possible if the fitness costs to a *D* allele (i.e., (1 − *zh*_*z*_)(1 − *g*)) are more than outweighed by an increase in fitness due to the realised killing advantage (i.e., (1 + *ae*)). In particular, invasion is more likely the more recessive the cost *z* (decreasing *h*_*z*_, compare columns in Figure 3). In the absence of costs (*g* = 0, *z* = 0), or in the presence of only fully recessive diploid costs (*h*_*z*_ = 0), invasion is possible if *ae* > 0.

In the absence of both costs (*g* = 0 and *z* = 0 or *h*_*z*_ = 0) and killing advantage (*a* = 0), *L*_*D*_ = 1, resulting in equality in condition (10). In this case, a second-order stability analysis (Otto and Day, 2011, p. 166) shows that invasion is possible if *e* > 0. Thus, a simple spore killer without fitness costs and killing advantage can still invade. Under these conditions a spore killer is a purely relative drive, which makes it sensitive to extinction due to drift (see Section 3.3).

A linear stability analysis of the fixation equilibrium, 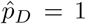, shows that it is stable with respect to invasion of the *d* allele when the fitness of *D* in a homozygote is higher than that of *d* in a heterozygote, *L*_*DD*_ > *L*_*d*_. Based on equations (2a) and (2c), this is the case when

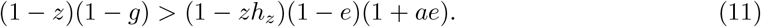

From this inequality we can see that fixation becomes less likely with increasing costs *g* and *z*. Furthermore, fixation becomes also less likely with a decreasing dominance parameter *h*_*z*_ = 0, increasing killing advantage *a* and decreasing killing efficiency *e*, which all favor the *d* allele when the frequency of *D* is high. Killing advantage and dominance level only matter if killing is incomplete (*e* < 1).

The two conditions for invasion and fixation lead to four possible outcomes. First, if *L*_*D*_ ≥ 1 and *L*_*DD*_ > *L*_*d*_, then the spore killer can invade and reach fixation, 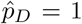 is a globally stable equilibrium. Second, if *L*_*D*_ < 1 and *L*_*DD*_ < *L*_*d*_, the spore killer cannot invade and 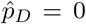 is a globally stable equilibrium. Third, if *L*_*D*_ ≥ 1 and *L*_*DD*_ < *L*_*d*_, the spore killer can invade and coexist with the sensitive allele at a stable polymorphism. Fourth, if *L*_*D*_ < 1 and *L*_*DD*_ > *L*_*d*_, the spore killer cannot invade from low frequencies, but it can reach fixation when starting from a frequency larger then the equilibrium value given by equation (9).

The conditions for stable coexistence, *L*_*D*_ ≥ 1 and *L*_*DD*_ < *L*_*d*_, impose the following requirements on the model parameters. First, killing has to be incomplete (*e* < 1), else fixation of the killer cannot be stopped whenever invasion is possible. Second, some fitness costs have to exist (*z* > 0 or *g* > 0), since otherwise with *e* > 0 the condition for fixation would always be fulfilled (recall that *a* < 1). Furthermore, from inequalities (10) and (11) we can see that a higher recessivity of the diploid costs, i.e., lower *h*_*z*_, favors the invasion of *D* but disfavors its fixation, and thus facilitates coexistence (Figure 3). Stable coexistence is also possible under an additive diploid fitness cost provided that the killing advantage *a* is sufficiently large (purple regions in the second column in Figure 3). This is possible because the fitness of the *d* allele, *L*_*d*_, represented by the right-hand side of inequality (11), increases with increasing killing advantage as this benefits *d* spores that survive killing.

In summary, in the *Neurospora* model: (i) A spore killer can invade if it bears no fitness costs, or if the costs are out-weighed by a killing advantage. (ii) If the killer allele can invade, then stable coexistence with the sensitive allele is possible provided that killing is incomplete (*e* < 1) and there are either sufficiently strong fitness costs to the spore killer or a strong killing advantage that benefits the sensitive allele in spores surviving killing.

#### 3.1.2 Invasion probability

Our stochastic simulations show that invasion of the killing allele *D* is possible whenever the equilibrium 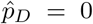 is unstable. For diploid costs this is shown in Figure 4 (see Figure S2 for haploid costs). The probability of invasion increases as costs and their dominance decrease. Furthermore, when the spore killer is associated with a killing advantage, fitness costs or both, then the dependency of the invasion probability on population size becomes negligible (see Figure S20). The special case with neither fitness costs nor killing advantage is treated in Section 3.3.

**Figure 4:**
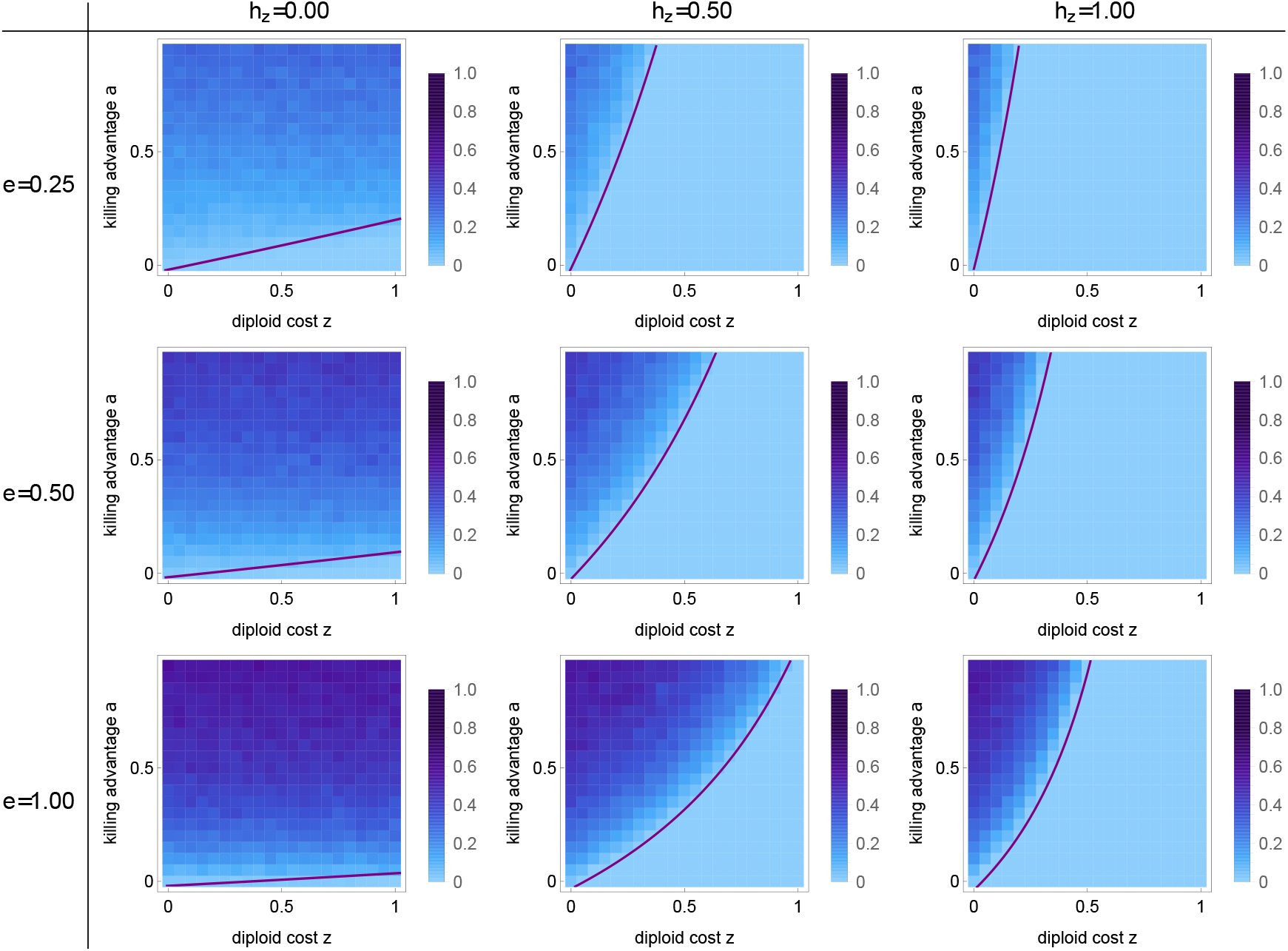
Invasion probability of a spore killer in the *Neurospora* model with diploid fitness costs *z*. Parameters are the fitness costs *z* and their dominance *h*_*z*_, the killing advantage *a*, and the killing efficiency *e*. Each panel consists of 21 × 21 parameter combinations and shades of blue indicate the invasion probability estimated from 10^3^ simulations of a Wright-Fisher process with population size 1000. Parameter combinations above and to the left of the purple line allow for invasion of the spore killer according to the deterministic model. Other parameters as in Figure 3.

### 3.2 Podospora anserina

#### 3.2.1 Deterministic model

The dynamics of the spore-killing allele *D* are affected by all parameters listed in the legend of Figure 2 and a complete analysis of all parameter combinations is beyond our scope. In the following, we analyze this model in two steps. In the first step we focus on the simplified *Podospora* model described in Section 2.2.2. This allows us to analytically investigate the effects of first- and second-division segregation (FDS and SDS, respectively) and of the dikaryotic costs *k*. In the second step, we a analyze the full model that additionally includes selfing, which occurs with probability *s*, monokaryons and their associated costs *k*_*m*_, and costs due to inbreeding *i*. For this second step we rely on numerical parameter sweeps. Table 1 summarizes the parameter combinations we examine and the corresponding Figures.

**Table 1:**
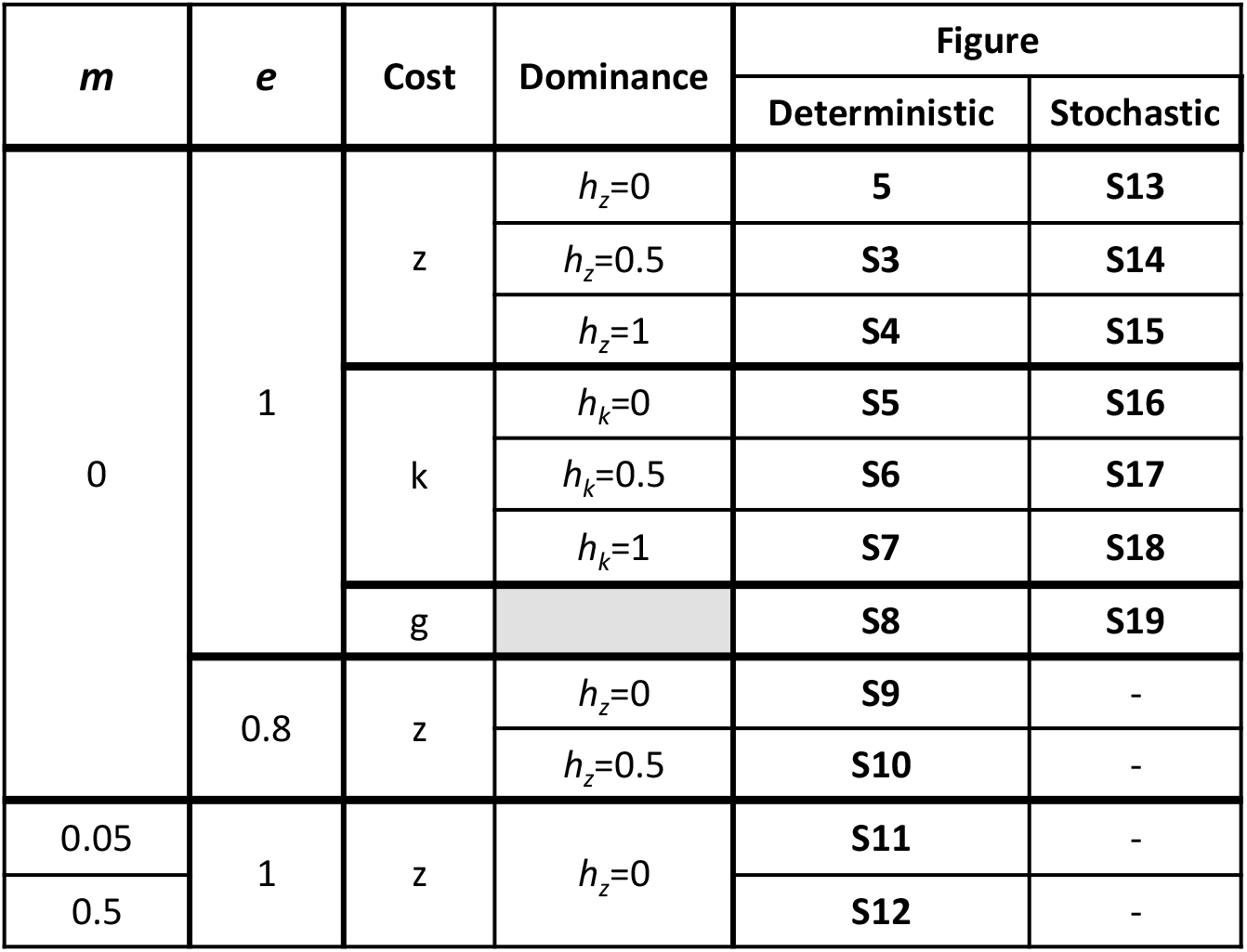
Overview of the parameter combinations investigated in the *Podospora* model.

#### 3.2.2. The simplified *Podospora* model

Since the simplified *Podospora* model follows the same recursion equation as the *Neurospora* model (equation (1)) but with modified fitness values (compare equations (3) and (2)) the two models have the same equilibrium structure. The first column in Figure 5 shows how the equilibria are affected by the killing advantage *a*, the probability of FDS *f* and a recessive diploid fitness cost *z*. Results for other levels of dominance, dikaryotic cost *k* at different levels of dominance and gametic cost *g* are shown in the first column of the corresponding Figures as indicated in Table 1.

**Figure 5:**
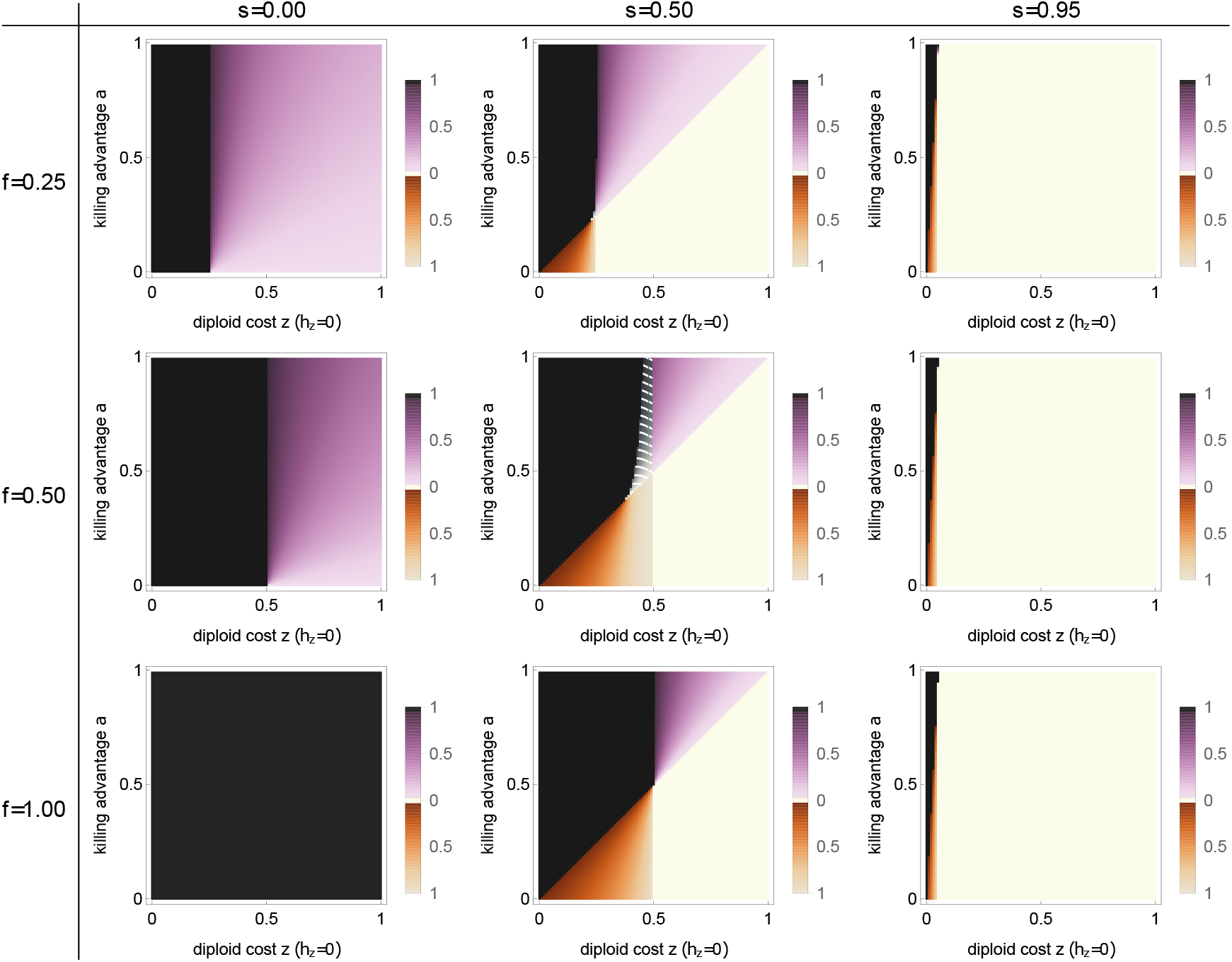
Bifurcation analysis of the *Podospora* model with recessive (*h*_*z*_ = 0) diploid fitness costs *z*. The bifurcation parameters are diploid fitness costs *z*, killing advantage *a*, selfing rate *s*, and probability of first-division segregation *f*. Extinction 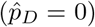 and fixation 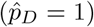 of the killer allele *D* always constitute equilibria. Additionally, one or two interior equilibria are possible. Parameter regions are color-coded as follows: **white**, *D* cannot invade and 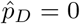 is a globally stable equilibrium; **black**, *D* can invade and reach fixation, 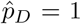 is a globally stable equilibrium; **purple**, *D* can invade but cannot reach fixation and coexists with the non-killing allele *d* at a globally stable interior equilibrium 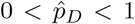 whose value is given by the shade of purple; **brown**, the two boundary equilibria 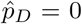 and 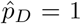 are stable, with their basins of attraction separated by an unstable interior equilibrium 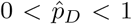 6 whose value is given by the shade of brown; **gray with white stripes**, two interior equilibria exist, the equilibrium with the lower value being stable, meaning that *D* can invade and coexist with *d* at a stable interior equilibrium, whose value is given by the shade of gray. Each panel is based on 100 × 100 parameter combinations. Other fixed parameters: *e* = 1, *m* = 0, other fitness costs are zero.

The killing allele *D* can invade if *L*_*D*_ > 1, which is the case when

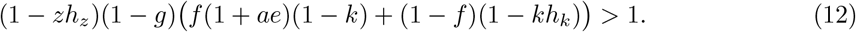

The left-hand side describes the fitness of the killer allele *D* when occurring in heterozygotes, and as long as the *D* allele is rare it indeed only occurs in heterozygous zygotes resulting from random mating. This explains why diploid costs *z* only occur in the factor (1 − *zh*_*z*_), the reduction in fitness of diploids carrying one copy of the *D* allele. Furthermore, the gametic cost *g* affects all *D* alleles in the same manner, independent of whether they derive from FDS or SDS, and therefore reduces fitness by a common factor (1 − *g*).

The situation is more complicated for the dikaryotic cost *k*. FDS occurring in heterozygous zygotes produces *D*:*D* dikaryons that experience a reduction of fitness by a factor (1 − *k*) while SDS produces *D*:*d* dikaryons that experience a reduction of fitness by a factor (1 − *kh*_*k*_). Thus, the *D* allele experiences the full cost *k* proportionally to the probability of FDS. As a result, under diploid and dikaryotic costs of equal magnitude and equal level of dominance, dikaryotic costs reduce the fitness of the killer allele more strongly than diploid costs and increasingly so with increased levels of FDS. This holds true whenever dikaryotic costs are not fully dominant (*h*_*k*_ < 1). If they are fully dominant, then (1 − *k*) can be factored out from the term in the third bracket on the left-hand side of inequality (13), which then shows that it has the same effect as a diploid or gametic cost. The effect of the dikaryotic compared to the diploid cost on the possibility for the *D* allele to invade can be seen when comparing Figure 5 with Figure S5 and Figure S3 with S6.

Whether or not FDS benefits allele *D* depends on a balance between two forces. On the one hand, FDS benefits *D* because it creates opportunities for killing and a killing advantage to arise (see Figure 2). On the other hand, SDS avoids the production of *D*:*D* dikaryons and can therefore benefit the *D* allele when dikaryotic costs *k* are high and recessive. More importantly, based on condition (12) it is easy to show that increased levels of FDS favor the *D* allele whenever its invasion is possible.

Finally, we highlight two special cases regarding the condition for invasion. First, with a fully recessive diploid fitness costs (*h*_*z*_ = 0) and in the absence of gametic and dikaryotic costs, invasion is always possible as long as *a* > 0 (see first columns in Figure 5). Second, the absence of both a killing advantage (*a* = 0) and costs (*z* = *k* = *g* = 0) represents the same special case of a purely relative drive described for the *Neurospora* model, for which invasion is possible as long as *e* > 0.

The *D* allele reaches fixation when *L*_*DD*_ > *L*_*d*_, which is the case when

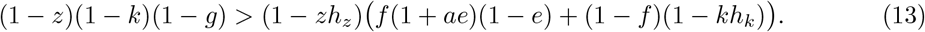

Whether or not FDS benefits fixation of the *D* allele depends on its effect on the *d* allele when rare (right-hand side of inequality (13)). When rare, the *d* allele occurs mostly in heterozygotes. With FDS, allele *d* pays a cost due to spore killing, which can be mitigated by a killing advantage if killing is incomplete. On the other hand, the fitness of the *d* allele following SDS is reduced by the factor (1 − *kh*_*k*_) due to being paired with the *D* allele in *D*:*d* dikaryons. By differentiating the right-hand side of inequality (13) with respect to *f* we find that FDS favors fixation of the *D* allele if

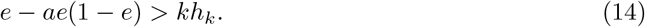

This condition is always fulfilled in the case of complete killing (*e* = 1), or fully recessive dikaryotic cost (*h*_*k*_ = 0) or absence of that cost (*k* = 0). This finding is confirmed by the first column of Figures 5, S3-S8, in which *e* = 1, and S9 and S10, in which *k* = 0.

One consequence of SDS (*f* < 1) is that coexistence of the alleles *D* and *d* becomes possible even in the presence of complete killing (*e* = 1), while in the Neurospora model complete killing always results in the fixation of the *D* allele whenever invasion is possible. This coexistence requires that the *D* allele suffers a recessive cost (*h*_*z*_, *h*_*k*_ < 1*/*2). For example, for the case of a fully recessive diploid cost (*h*_*z*_ = 0) and in the absence of other costs, the condition for invasion, inequality (12), simplifies to *fa* > 0 and the one for fixation, inequality (13), to *f* > *z*, meaning that coexistence is possible with *e* = 1 as long as *f* < *z* (see first column in Figure 5). Finally, we note that fully recessive dikaryotic costs (*h*_*k*_ = 0) affect the condition for fixation in the same manner as gametic costs (compare the first columns of Figures S5 and S8).

#### 3.2.3 The complete *Podospora* model

Compared to the simplified *Podospora* model, the full model includes the possibility for selfing and monokaryons and it is these two aspects on which we focus in this section. The full *Podospora* model allows for one additional outcome, namely, the killing allele *D* can invade and there are two internal equilibria, with the lower one stable and the upper one unstable. However, this case occurs only in a small region of the parameter space (Figure 5, dashed area).

##### (i) Effect of selfing

Here, we explore the effect of selfing on the dynamics of the spore killer allele *D*. We first focus on the case of complete killing (*e* = 1), no monokaryons (*m* = 0), and fully recessive diploid cost (*h*_*z*_=0), and subsequently relax these restrictions in sections (ii) and (iii). Given these restrictions, the second and third columns in Figure 5 show the effect of an intermediate and a high selfing probability, respectively. For these findings we can provide intuitive explanations but no analytical proof. The following observations can be made.

First, invasion of the *D* allele becomes more difficult with increasing selfing probability. For instance, for the parameters used in Figure 5 and with *s* = 0.5, invasion becomes impossible if the diploid cost exceeds the killing advantage (*z* > *a*), whereas invasion is always possible with *s* = 0 (compare first and second column in Figure 5). For higher selfing probabilities invasion is restricted to even smaller regions of the parameter space. The reason is that *D* : *D* dikaryons that reproduce by selfing result in *DD* diploids and these pay the full diploid fitness cost. Random mating instead produces mostly *Dd* diploids during invasion due to the low frequency of *D*. The reduction of the parameter space allowing for invasion of the *D* allele due to this mechanisms is more pronounced the more recessive the diploid costs (lower *h*_*z*_, compare the second column in Figures 5, S3 and S4).

Second, generally, fixation of the *D* allele becomes more difficult with increasing selfing probability. The reason is that selfing maintains *d*:*d* dikaryons and thereby protects the *d* allele from spore killing. However, this effect of selfing on the fixation of *D* becomes visible only at sufficiently low levels of SDS, because high levels of SDS also shield *d* from spore killing. For instance, Figure 5 shows that increasing the selfing probability from *s* = 0 to *s* = 0.5 does not affect fixation of the *D* allele at low (*f* = 0.25) or moderate (*f* = 0.5) levels of FDS but does hinder fixation at high levels (*f* = 1). Qualitatively similar observations hold for additive and dominant diploid costs (Figure S3 and S4). This effect is further modulated by the level of dominance of the dikaryotic costs. Theses costs decrease the effectiveness of SDS in protecting the *d* allele, which gives SDS a comparatively smaller role in preventing fixation of *D* with increasing *h*_*k*_ (compare Figure S5 with S6 and S7).

We note that selfing can involve a fitness cost *i* due to inbreeding at all loci, which applies to individuals regardless of their genotype. This cost reduces the effective selfing rate by removing a portion of selfing individuals and does not interact with the dynamics of the spore killer.

##### (ii) Effect of incomplete killing

We restrict ourselves to the case of diploid costs *z*, which are more conducive to invasion of the killing allele than dikaryotic and gametic costs. Figure S9 shows a bifurcation diagram analogous to Figure 5 but with *e* = 0.8. As expected, lowering the killing efficiency lowers the upper boundaries of the diploid costs below which invasion and fixation of the spore killer are possible. This observation also holds under additive costs (compare Figures S3 and S10). As can be seen, incomplete killing (*e* < 1) enables coexistence in the presence of an additive diploid fitness cost.

##### (iii) Effect of monokaryons

A final aspect of the *P. anserina* life cycle that we explore is the effect of monokaryons on the dynamics of the spore killer. As can be seen from Figure 2, monokaryons are not able to self and they allow for a small amount of spore killing even in the case of SDS. Thus, by limiting the selfing rate and by allowing killing even when the probability of FDS is low we expect that monokaryons facilitate both invasion and fixation of the spore killer.

For the case of recessive diploid costs (*h*_*z*_ = 0) and no other costs we can formalize the effect of monokaryons as follows. Based on the life cycle in Figure 2, we can express the proportion of killed spores, *K*, as

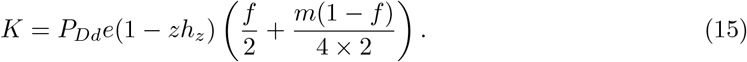

Thus, the number of killed spores increases with the product of the proportion of monokaryons and SDS events at meiosis, *m*(1 − *f*), and with the frequency *P*_*Dd*_ of heterozygotes in the population (which itself increases with *m* due to outcrossing).

For 5% and 50% of asci containing a pair of monokaryons, Figures S11 and S12, respectively, show how monokaryons increase the upper boundaries of the diploid costs below which invasion and fixation of the spore killer are possible. These figures should be compared to Figure 5, which corresponds to the same parameter values but without monokaryons. A frequency of 5% is in the range expected in natural populations, but we also consider 50% to magnify the effect of monokaryons. In the case of 50% of asci containing monokaryons, the expected effects of monokaryons become clearly visible (Figure S12). With 5% of asci containing monokaryons, the effect of monokaryons appears negligible, indicating that they probably do not greatly affect spore killer dynamics under natural conditions.

The results for the *Podospora* model can be summarized as follows. (i) A spore killer can invade if it bears no fitness costs, or if the costs are outweighed by the fitness benefit due to a killing advantage. (ii) Selfing makes invasion and fixation of the spore killer more difficult. This is because it reduces the frequency of heterozygotes and increases the chance that recessive costs become expressed. Selfing interacts with the probability of FDS to determine the effective killing rate. (iii) Coexistence between a spore killer and a sensitive allele is possible with incomplete killing efficiency (*e* < 1) in combination with recessive or additive fitness cost, and remains possible with fully efficient killing (*e* = 1) in combination with a recessive fitness cost and either some amount of selfing or SDS (*f* < 1).

#### 3.2.4 Invasion probability

Our stochastic simulations confirm that invasion of the killing allele *D* is possible whenever the equilibrium 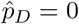 is unstable. The probability of invasion increases with the probability of first-division segregation *f* and the killing advantage *a*, which both increase the selective advantage of the spore killer. In turn, invasion probability decreases with fitness costs. We refer the reader to Table 1 for an overview of the parameter combinations investigated and the corresponding figures.

### 3.3 Invasion probability of a spore killer without killing advantage

A spore killer without killing advantage is a purely relative drive, and consequently its invasion probability tends to zero as the population size *N* becomes large (Nauta and Hoekstra, 1993). However, it has not been investigated how rapidly this invasion probability declines as *N* grows, and how the contrasting effects of population size on invasion probability and mutational supply interact to determine the overall rate of invasion of spore killers.

#### 3.3.1 Invasion probability as a function of population size *N*

We study the simplest possible model of a newly-arisen spore killer with complete killing (*e* = 1), assuming random mating (no selfing). Employing a heuristic method from Desai and Fisher (2007, p. 1763), we show in Appendix 1 that, once the spore killer exceeds 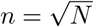 copies, the expected increase in its copy number over the subsequent *n* generations (∼*n*^3^*/N*) is greater than typical decreases in copy number by random drift over the same period (∼*n*), so that the spore killer has very likely escaped stochastic loss. The dynamics of the spore killer before reaching 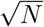 copies are dominated by random drift, so the probability that it attains the required 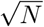 copies—i.e., its invasion probability—is approximately 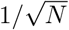 (Crow and Kimura, 1970). This approximation agrees well with estimates obtained from simulations (Figure 6a).

**Figure 6:**
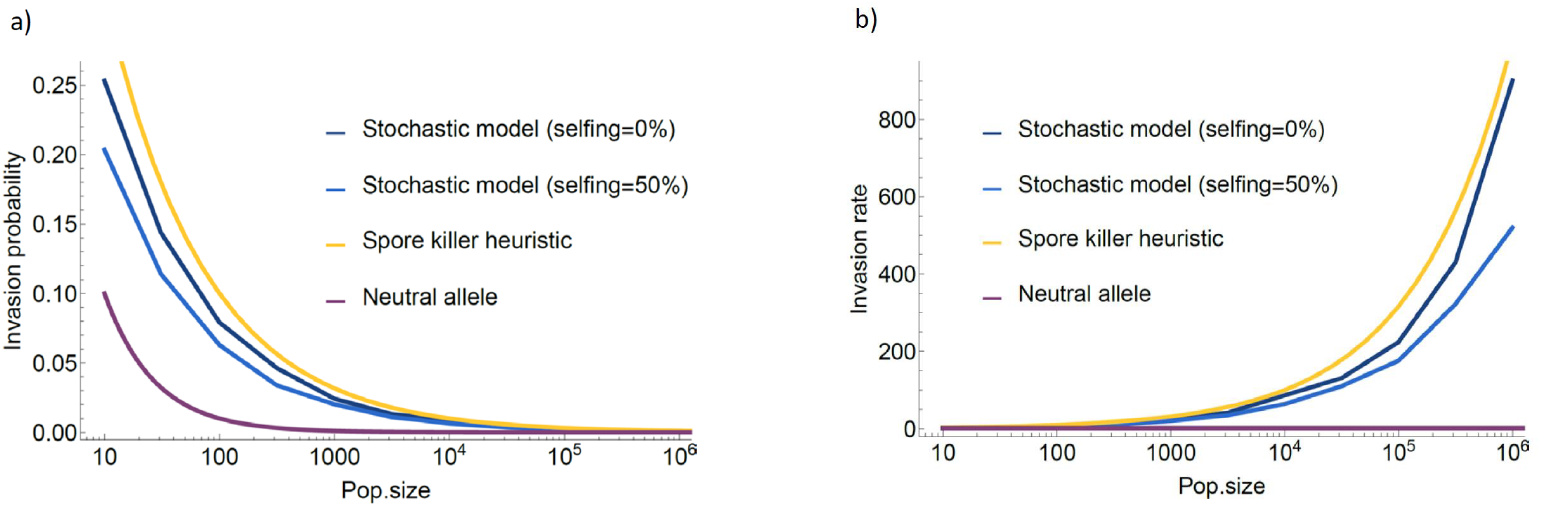
Invasion probability (a) and invasion rate (b) of a spore killer as a function of population size. Fitness costs or killing advantage are absent. A comparison is made between the approximation developed in Section 3.3 (spore killer heuristic), the stochastic invasion model, and the stochastic invasion model with 50% selfing rate. A neutral allele is also represented for reference.

#### 3.3.2 Invasion rate as a function of population size *N*

If spore killers arise at a rate *µ* per replication, then their rate of appearance per generation is *Nµ*. Therefore, the per-generation rate of invasion of such spore killers is 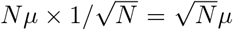. The invasion rate is thus an increasing function of population size (Figure 6b), despite the fact that the invasion probability of each individual spore killer decreases as *N* increases, tending to zero in the large-population limit. Additional life-history features, such as selfing, affect the invasion rate but do not affect the positive scaling with population size (Figure 6b).

#### 3.3.3 Invasion probability and population structure

The dependence of the invasion probability on population size suggests that population structure could significantly affect the rate at which spore killers invade. Suppose that the population of size *N* is subdivided into *M* demes, each of size *m* = *N/M*. The overall arrival rate of new spore killers is unchanged (*Nµ*), but, since invasion of one deme guarantees population-wide invasion (unless *m* is very small), the invasion probability is 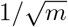. The invasion rate is therefore 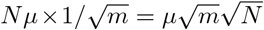, which is larger than in an unstructured population by a factor of 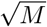. A similar result has been obtained for recessive beneficial mutations (Gale, 1990, p. 180-181), which are under similar frequency-dependent selection (see Appendix 1).

## 4 Discussion

Our study is novel in exploring the effect of several aspects of fungal life cycles on the conditions under which a spore-killing allele can invade and subsequently stably coexist with a non-killing allele. In ascomycete fungi, spore killing takes place within the ascus, and our model is based on a detailed mechanistic understanding of ascus composition (see Figure 2). Our results show that following the possible compositions of spores within an ascus is necessary to fully understand the dynamics of a spore-killing allele. Another novel aspect of our study is the development of stochastic models to investigate the invasion probability of a spore-killing allele, complementing our deterministic analysis. Our model is based on a single allele responsible for both spore killing and resistance to spore killing. Thus, there is no recombination between the two functions. This feature is consistent with the picture emerging from recent genetic characterization of spore killers in several species of ascomycete fungi (Nuckolls et al., 2017; Hu et al., 2017; Vogan et al., 2019; Svedberg et al., 2020). The only other theoretical study of spore killing we are aware of (Nauta and Hoekstra, 1993) focuses on the role of recombination between the killing and resistance functions, which was appropriate given what was known about the genetic architecture of spore killers at that time in *Neurospora*, but appears now to be the exception rather than the rule.

In the following, we first discuss the general insights that our model reveals about the dynamics of spore killers. We then discuss our results in the light of data from natural systems, with particular attention paid to the spore killer systems *Spk-1* in *Neurospora* and *Spok* in *Podopsora*, which inspired our model. Throughout, we compare our findings to theoretical and empirical results from male and female meiotic drivers in animals and plants.

### 4.1 Spore killers in theory

#### 4.1.1 Selective advantage and invasion of a spore killer

We start by comparing the dynamics of spore killers in our model with Hartl’s (1972) general model of sperm and pollen killers. In Hartl’s model, gamete killers kill in the homozygote form, which could be seen as “self-killing” or “suicide” as effectively the killer allele eliminates other copies of itself. This “self-killing” does not provide any fitness advantage for the killer in the absence of a killing advantage. In contrast, spore killers do not “self-kill”, which allows them to obtain a positively frequency-dependent selective advantage simply by killing. In agreement with Nauta and Hoekstra (1993), we find that this frequency-dependent selective advantage tends to zero when the frequency of the spore killer is close to zero. Thus, in the absence of any killing advantage, the selective advantage of a spore killer is minimal at the onset of invasion (except in small populations), and so invasion of spore killers can be prevented by any non-recessive fitness costs, or simply by stochastic loss (see Section 3.3). This distinguishes spore killers from female drive systems, in which the selective advantage induced by drive itself can be substantial at all frequencies of the driver, allowing female drivers to invade even when they are associated with substantial fitness costs (e.g. Hall and Dawe, 2018).

Although the invasion probability of a spore killer without killing advantage is substantially lower than that of a female driver, it is not negligible (Section 3.3). Moreover, owing to the frequency-dependent nature of the spore killer’s selective advantage, small or fragmented populations may represent easier targets for invasion. Thus, the study of spore killers in structured populations, which we have only briefly addressed in Section 3.3, merits further study. In addition, our calculations show that, even in the absence of a killing advantage, the invasion rate of spore killers increases with population size, as the increase in mutational supply of spore killers more than compensates for the decrease in their individual invasion probabilities (Section 3.3). This last point depends, of course, on the mechanism of origin of spore killers, which we discuss in the next section.

In addition to the action of killing itself, we propose that spore killers could induce a killing advantage, i.e., a net fitness benefit from killing, either in the form of compensation or reduced local competition for resources. In the model of Hartl (1972), compensation plays a crucial role for gamete killers, granting them a selective advantage. Although we cannot draw a direct parallel to Hartl’s model, which focuses on the role of gamete number to fecundity functions, we also find that a killing advantage is crucial for the invasion of a spore killer, conferring on it a frequency-independent advantage that, being present even when the spore killer is rare, can therefore promote its initial invasion. In particular, a killing advantage reduces the chance of stochastic loss of a killing allele. It is therefore vital, in future empirical studies of spore killer systems, to determine whether and to what extent a killing advantage is present.

#### 4.1.2 Coexistence of a spore-killing allele with a non-killing allele

We also investigate the conditions for stable coexistence of the killing and non-killing alleles. Meiotic drivers (MDs) are expected often to spread rapidly to fixation (Lindholm et al., 2016), rather than coexisting with their non-driver alleles in a stable polymorphism—and when a MD is fixed, drive does not occur, rendering the MD undetectable. The MDs that are observed in natural conditions are therefore expected to exhibit (possibly unusual) properties that promote their maintainance in stable polymorphisms. Understanding these properties is therefore an important task of theoretical studies of meiotic drive. Classically, in male and female drive, recessive fitness costs associated with the driving allele are required for coexistence (e.g., Hartl, 1970; Fishman and Kelly, 2015; Lewontin and Dunn, 1960; Holman et al., 2015)—invasion is permitted because the costs are not expressed in heterozygotes, while fixation is prevented because the costs are expressed in homozygotes. We find this dynamic in our models for both *Neurospora* and *Podospora*. Fitness costs are needed for coexistence to be possible and recessive costs increase the parameter space that supports coexistence (Figure 5). For coexistence to be possible, it is also necessary that killing is incomplete, so that the sensitive allele obtains a fitness benefit when in a diploid heterozygote. Incomplete killing can result from the killing efficiency being less than 100% and, in the case of *Podospora*, from second-division segregation.

Intriguingly, we find that coexistence is also possible even if fitness costs to the spore killer are not recessive. Such coexistence occurs because sensitive spores that survive killing also benefit from the killing advantage. We believe that this assumption is empirically reasonable given the two possible scenarios that we envision underlying a killing advantage, namely compensation and reduced local competition. Compensation results in additional spores—both killer and sensitive— being produced by the parent in response to killing. If a killing advantage occurs via reduced local competition with siblings, then the same reasoning applies and both types of surviving spores obtain a fitness advantage. Coexistence can occur in this case because the benefit to the surviving sensitive spores prevents fixation of the spore killer by raising the fitness of the sensitive allele in heterozygotes above the fitness of the spore killer in homozygotes. Thus, incomplete killing can result in coexistence just as recessive costs can (or overdominance, Hartl, 1970), but the underlying biology is distinct.

#### 4.1.3 Mating system and spore killer dynamics

We find that the rate of selfing of the host has a negative effect on the invasion of a spore killer. The reason is that selfing decreases the frequency of heterozygotes (in which spore killing occurs), and magnifies potential fitness costs, generally slowing down invasion. Because of this latter point, our model predicts that a spore killer can invade a population with a high selfing rate only when it is associated with low fitness costs. Moreover, coexistence is then unlikely. We expect inbreeding to have the same effect as selfing, and analyzing spore killer models in which the assumption of random mating is relaxed could be an interesting next step. This influence of selfing further suggests that mating behavior itself could evolve as a defence against spore killers, as previously suggested by Lewontin and Dunn (1960). Along the same lines, Bull (2017) and Bull et al. (2019) have developed models showing that inbreeding can evolve as an efficient response to costly meiotic drivers, while Burt and Trivers (1998) have shown that obligatorily outcrossing plant species are more susceptible to costly selfish genetic elements.

### 4.2 Insights from natural systems

#### 4.2.1 How much do we know about spore killers in nature?

In several model systems of male and female drive, the molecular mechanism of the MD, its fitness effects, and the biology of the host are known to a sufficient extent that population genetics models can predict the frequency of the MD in natural or laboratory populations with impressive accuracy (e.g. Lewontin and Dunn, 1960; Fishman and Kelly, 2015). In the case of spore killers, however, although several recent publications have shed light on the genetic and molecular basis of their driving action (Hu et al., 2017; Nuckolls et al., 2017; Vogan et al., 2019; Svedberg et al., 2020), many unknowns remain, particularly regarding the ecology of the hosts. In this section, we summarize the available knowledge and use it to put our results in perspective and to suggest future directions for empirical research.

We focus on the three spore killers in fungal hosts that are best understood: the *Spok* gene family in *Podospora anserina* (Vogan et al., 2019) and *Spk-1* in *Neurospora sitophila* (Svedberg et al., 2020)—both of which directly inspired our models—and finally the *wtf* gene family in *Schizosaccharomyces pombe* (Hu et al., 2017; Nuckolls et al., 2017), which is also well studied and has many similarities with the first two. In all three cases, spore killing and resistance are governed by a single locus, as we have assumed in our model. Although we restrict ourselves to these three best-known example from model species, we believe that spore killers could be widespread in the fungal kingdom, or at the very least in ascomycetes where they have been observed in several taxa (Raju, 1994). Fungi show extreme diversity in their mode of reproduction (Nieuwenhuis and James, 2016), and our results suggest that high selfing rates may be a major factor in hampering the invasion of spore killers. Reproductive strategy may therefore be a key determinant for the presence of spore killers in a given species. Most fungi are able to reproduce sexually (Nieuwenhuis and James, 2016), and are therefore susceptible to spore killers in theory, but many have high rates of selfing (Nieuwenhuis and James, 2016), which our findings suggest could protect them efficiently from the invasion of spore killers.

Little is known about the origin of spore killers. It has been proposed that spore killing systems may arise neutrally in populations in which resistance to killing has already been fixed (Sweigart et al., 2019). According to this view, spore killers would then act as a strong type of hybrid incompatibility between diverging populations. However, our work shows that an active spore killer has a substantially greater chance of invading than a neutral allele, which suggests that selfish evolution of spore killers could be more likely. The *Spok*s, *Spk-1*, and *wtf* s all belong to large families of genes that occur across complexes of closely related fungal taxa. This suggests the possibility of horizontal gene transfer across species. For example, there is evidence that *Spk-1* in *N. sitophila* may have introgressed from the closely related *N. hispaniola* (Svedberg et al., 2020). In addition to their apparently frequent movements, *Spok* and *wtf* genes mutate rapidly (Hu et al., 2017; Nuckolls et al., 2017; Vogan et al., 2019), which could be the key to their success (see Section 3.3 for the importance of mutation rate).

#### 4.2.2 Insights from our models on spore killer dynamics in natural populations

The *Spok* gene family has several members present in the genomes of species of the *Podospora* genus. In *P. anserina* in particular, three genes are known: *Spok2, Spok3*, and *Spok4*. Any given individual of *P. anserina* might carry none, one, two, or all three *Spok* genes. With very stable recombination rates, the FDS probability that ultimately determines the killing rate of the *Spok* genes is determined by their position along the chromosomes. More than one copy of a *Spok* gene might occur in a single genome, but this seems to be very rare (Vogan et al., 2019). *Spok3* and *Spok4* occur in a genomic region known as the ‘*Spok* block’ (Vogan et al., 2019). The different *Spok* genes act independently, that is, carrying one of them does not protect against another one (Grognet et al., 2014; Vogan et al., 2019). Figure S21 shows the frequency of the three *Spok* genes in samples from a population of *P. anserina* near Wageningen in the Netherlands over a 17 year period (generation time can be as short as 11 days). None of the spore killers reached fixation or went extinct during this period. However, *Spok2* appears close to fixation while *Spok3* and *Spok4* occur at lower frequencies. Additionally, all individuals without *Spok2* seem to derive from a single deletion, thus, *Spok2* may have been fixed prior to 1993.

We can try to understand, with the help of predictions from our model, what characteristics of the different *Spok* genes may be responsible for their respective frequencies in the Wageningen population. We know that *Spok2* does not kill with full efficiency and has a FDS probability of about 40%, while *Spok3* and *Spok4* have a FDS probability ranging between 50% and 90%, depending on their position in the genome (Vogan et al., 2019). Yet, *Spok2* appears closer to fixation. If we take into account the presumably high selfing rate of *P. anserina* this observation is in agreement with our model, which predicts that FDS probability should matter little to the dynamic of the killer if selfing rate is high (outcrossing rate becomes limiting: compare the second and third columns in Figures 5 and S9). Our model also predicts that a high selfing rate combined with fitness costs severely reduces the scope for invasion of a spore killer (e.g. Figure S19). Thus, in this context, the fate of different *Spok* genes should be largely determined by their fitness cost but not by their FDS probability. And indeed, we know that *Spok3* and *Spok4* co-occur in the ‘*Spok* block’ (Vogan et al., 2019), which is known to be associated with fitness costs to its host (Vogan et al., 2020). This observation also confirms that increasing selfing rate may be a good defense mechanism against the invasion of spore killers, but that in that context the probability of FDS, regulated by the position of the *Spok* genes on the chromosomes, matters little in *P. anserina*.

The gene *Spk-1* in *N. sitophila* shows variation across populations, being respectively fixed and absent in two clades that coexist in sympatry, and polymorphic in a third clade where a form of resistance to the killer has evolved (Svedberg et al., 2020). This suggests that the dynamics of the same spore killer can vary in different populations. In the polymorphic clade, resistance has evolved in the form of reduced killing efficiency, leading to what appears to be coexistence (Svedberg et al., 2020).

Very little is known about the frequency of the *wtf* allele in natural populations of *S. pombe*. Killing efficiency is lower than 100% (Nuckolls et al., 2017; Núñez et al., 2020) and selfing is likely common in *S. pombe* (Nieuwenhuis and James, 2016; Tusso et al., 2019). Furthermore, *S. pombe* is able to perform haploid selfing, a feature that is not found in *P. anserina* and that we therefore did not incorporate in our model. Based on this information, we predict that the fitness costs of *wtf* must be low to allow for invasion, and that invading spore killers progress slowly and are sensitive to stochastic loss.

At least one suppressor counteracting the action of the spore killer has evolved in *S. pombe* (Nuckolls et al., 2017; Núñez et al., 2020). Suppressor genes are likely to evolve if given enough time (slow invasion or stable coexistence) and are found in many other drive systems, but not in *P. anserina* so far (Vogan et al., 2019).

The spore killers *Spk-1, Spok* and *wtf* are all small genomic regions without inversions, suggesting that they are not necessarily associated with hitchhiking deleterious mutations (but see Vogan et al., 2020). At the same time, all spore killers function with a poison-antidote mechanism targeting spores, and it is easy to envision direct fitness costs for production and exposure of spores or even vegetative mycelium to the toxin. These costs could be recessive if subject to a threshold dosage effect. To date, there is no definitive evidence for or against fitness costs of carrying these spore killers, except for *Spok3* and *Spok4*, which are contained within the ‘*Spok* block’ element. In the latter case, it is not clear whether the cost originates from the *Spok* genes themselves or other features of the block. Finally, there is evidence from laboratory studies for a killing advantage in *P. anserina* through a recovery in the number of spores produced when killing occurs (Vogan et al., 2020). However, more work is required to understand its importance in a natural setting.

### 4.3 Conclusions

Despite their particularities, we predict that spore killers should show similarities with well-studied systems of meiotic drive. For example, we expect fitness costs—likely but not necessarily recessive—to explain coexistence. Our study identifies characteristics of the MD loci that interact with the ecology and the life cycles of ascomycete fungi and are of importance for the dynamics of spore killers. These are: fitness costs, killing advantage, host population size, and the mating system. Although we find that a spore killer without costs or killing advantage is substantially more likely to invade than a neutral allele, killing advantage makes invasion substantially more likely. In contrast, selfing of the host and fitness costs associated with the killer can impede its invasion or halt its spread at intermediate frequencies.

With this work, we have explored the dynamics of spore killers in two representative species of ascomycete fungi, and have revealed novel aspects of the dynamics of these selfish genetic elements. Our work has revealed that many unknowns remain from both theoretical and empirical angles that prevent, at this stage, accuarte empirical prediction of spore killer dynamics. With the advent of artificial meiotic drives, a new world of possibilities opens for biological control (Esvelt et al., 2014). If spore killers are to be used for the control of fungal pest species, there is still much work that needs to be done in order to fully account for their dynamics. We suggest several points of focus for future research. Important empirical tasks will be to better understand the ecology of fungal hosts, in particular regarding their mating systems, as well as to better characterize interactions between spore killers and their hosts (fitness effects, killing advantage). From a theoretical perspective, we suggest that the role of population structure and the possibility for the evolution of suppressor genes in spore killers should be important aspects.

## Supplementary Material

**Figure S1:**
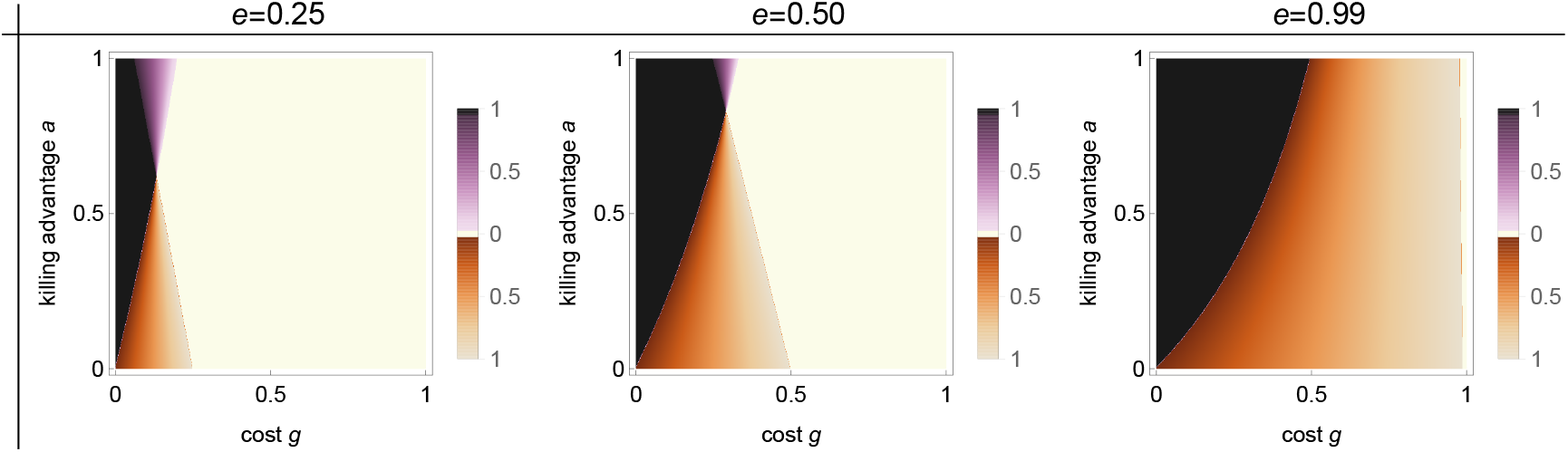
Bifurcation analysis of the *Neurospora* model with haploid fitness costs *g*. The bifurcation parameters are haploid fitness costs *g*, killing advantage *a*, and killing efficiency *e*. Extinction 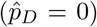 and fixation 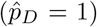 of the killer allele *D* always constitute equilibria. Additionally, one interior equilibrium is possible. Parameter regions are color coded as follows: **white**, *D* cannot invade and 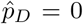 is a globally stable equilibrium; **black**, *D* can invade and reaches fixation, 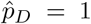 is a globally stable equilibrium; **purple**, *D* can invade but cannot reach fixation and coexists with the non-killing allele *d* at a globally stable interior equilibrium 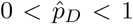, whose value is given by the shade of purple; **brown**, the two boundary equilibria 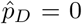 and 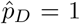 are stable, with their basins of attraction separated by an unstable interior equilibrium 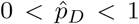 whose value is given by the shade of brown. Each panel is based on 1000×1000 parameter combinations. Fitness costs *z* = 0.

**Figure S2:**
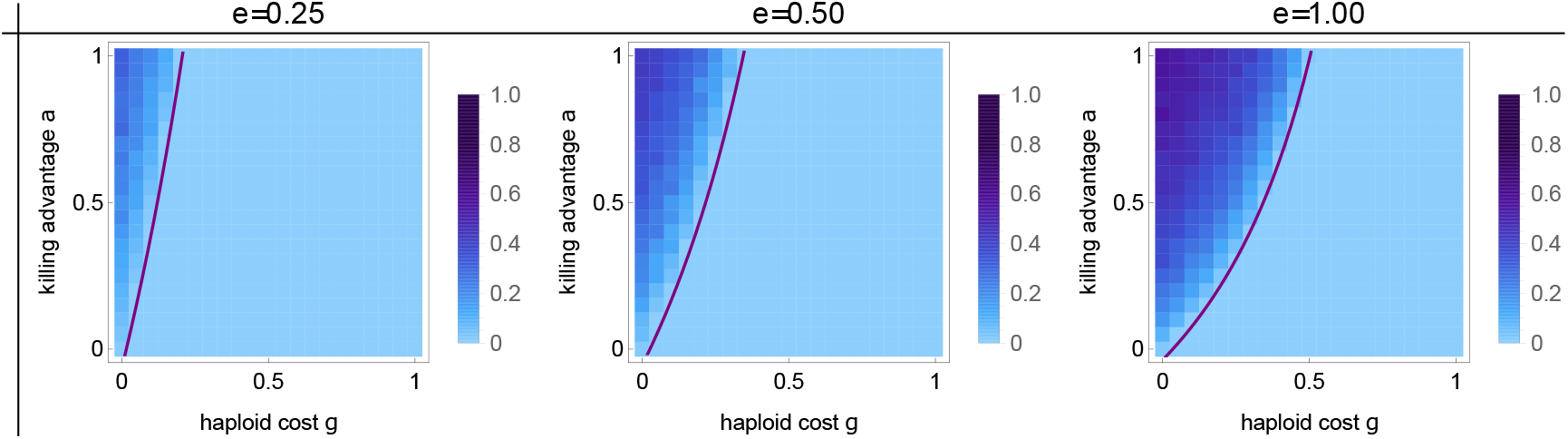
Invasion probability of a spore killer for the *Neurospora* model with haploid fitness costs *g*. Parameters are the fitness costs *g*, the killing advantage *a*, and the killing efficiency *e*. Each panel consists of 21×21 parameter combinations and shades of blue indicate the invasion probability estimated from 10^3^ stochastic Wright-Fisher simulation runs with a population size of 1000. Parameter combinations to the left and above the purple line allows for invasion of the spore killer according to the deterministic model. Other parameters as in Figure S1.

**Figure S3:**
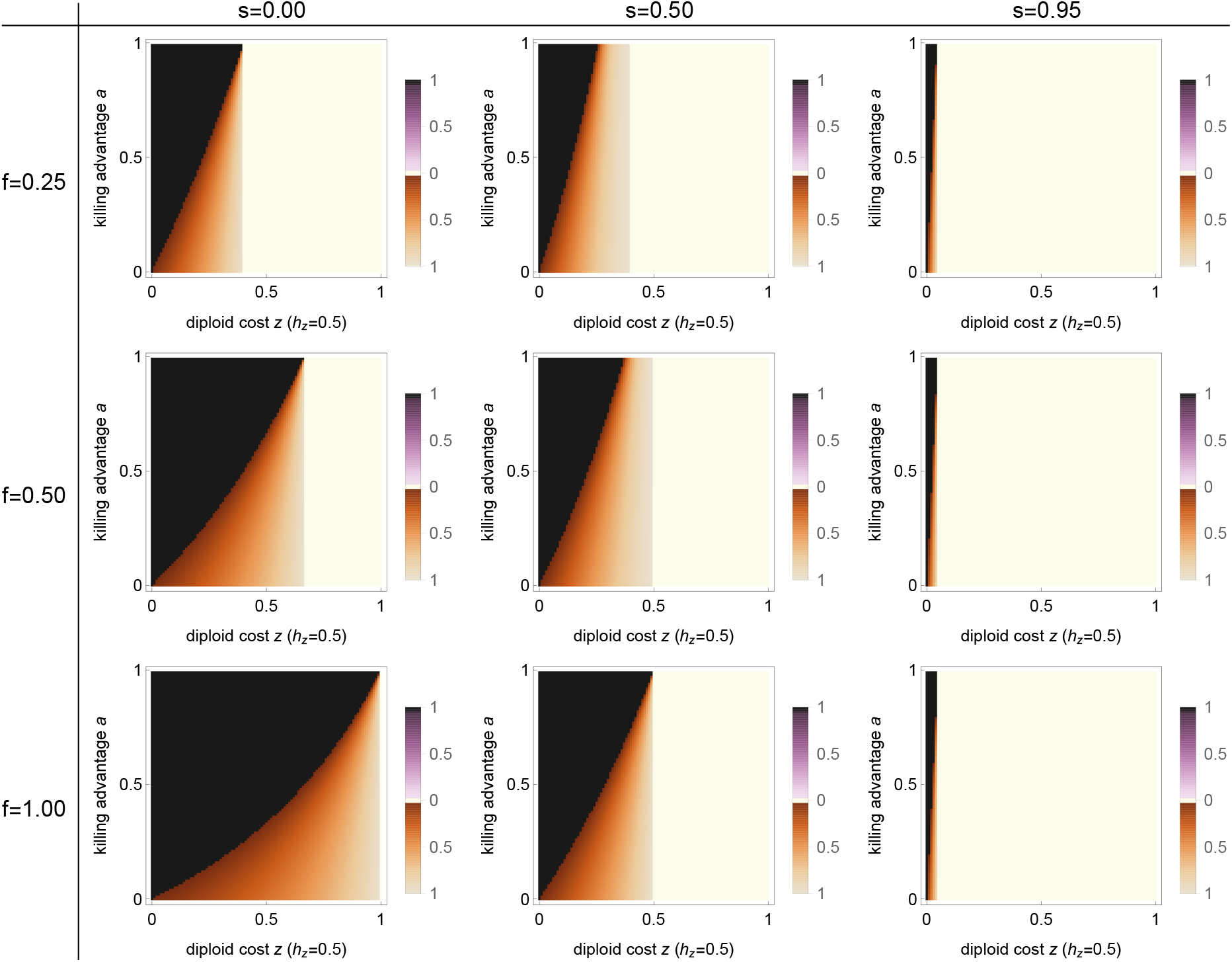
Bifurcation analysis of the *Podospora* model with additive (*h*_*z*_ = 0.5) diploid fitness costs *z*. The bifurcation parameters are diploid fitness costs *z*, killing advantage *a*, selfing rate *s* and probability of first-division segregation *f*. Extinction 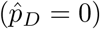 and fixation 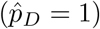 of the killer allele *D* always constitute equilibria. Additionally, one interior equilibrium is possible. Parameter regions are color coded as follows: **white**, *D* cannot invade and 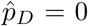 is a globally stable equilibrium; **black**, *D* can invade and reach fixation, 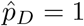 is a globally stable equilibrium; **purple**, *D* can invade but cannot reach fixation and coexists with the non-killing allele *d* at a globally stable interior equilibrium 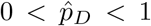, whose value is given by the shade of purple; **brown**, the two boundary equilibria 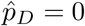 and 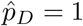 are stable, with their basins of attraction separated by an unstable interior equilibrium 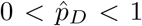 whose value is given by the shade of brown. Each panel is based on 100×100 parameter combinations. Other fixed parameters: *e* = 1, *m* = 0, other fitness costs equal to zero.

**Figure S4:**
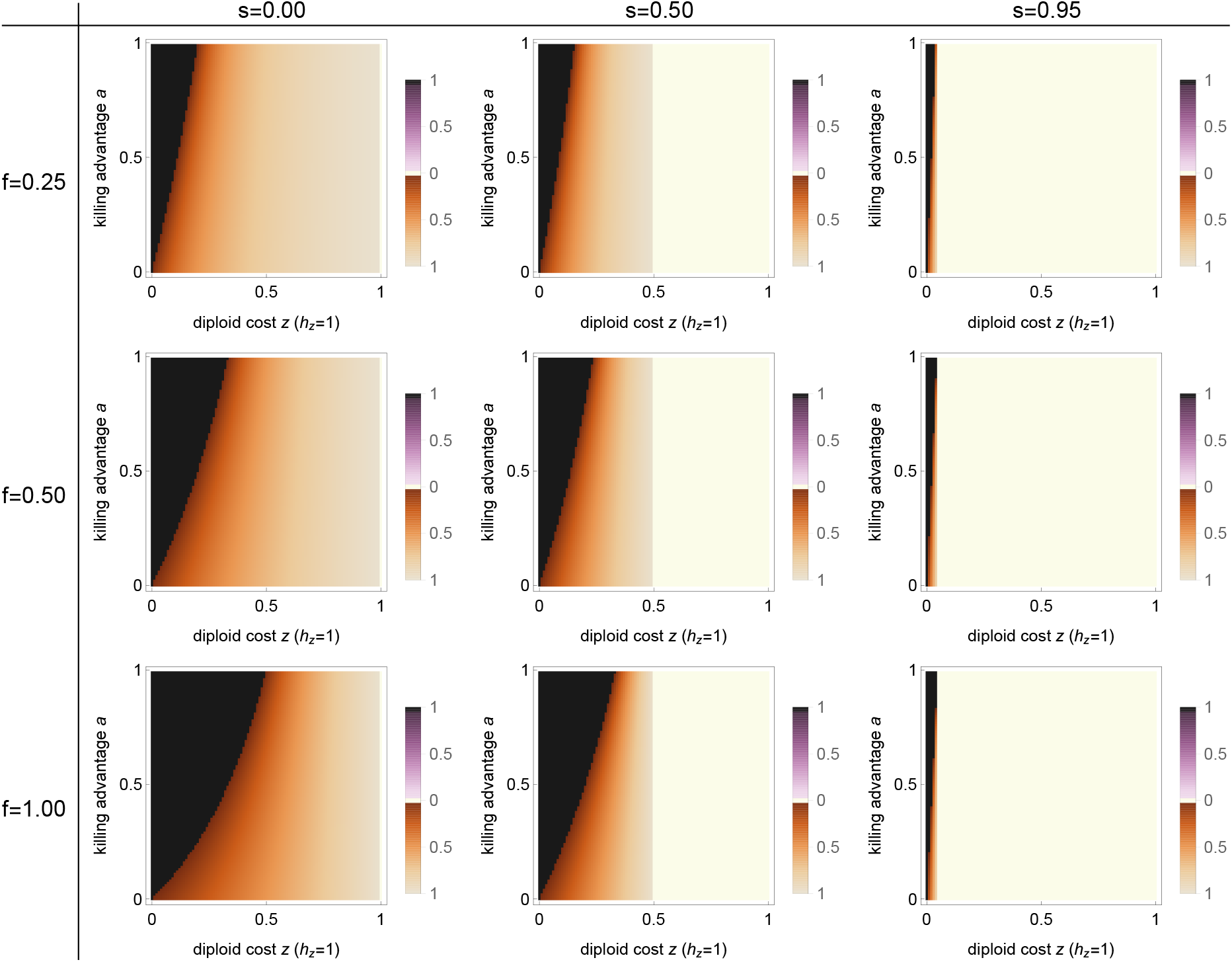
Bifurcation analysis of the *Podospora* model with dominant (*h*_*z*_ = 1) diploid fitness costs *z*. The bifurcation parameters are diploid fitness costs *z*, killing advantage *a*, selfing rate *s* and probability of first-division segregation *f*. Extinction 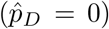 and fixation 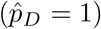 of the killer allele *D* always constitute equilibria. Additionally, one interior equilibrium is possible. Parameter regions are color coded as follows: **white**, *D* cannot invade and 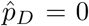 is a globally stable equilibrium; **black**, *D* can invade and reaches fixation, 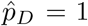 is a globally stable equilibrium; **purple**, *D* can invade but cannot reach fixation and coexists with the non-killing allele *d* at a globally stable interior equilibrium 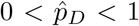, whose value is given by the shade of purple; **brown**, the two boundary equilibria 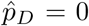 and 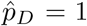 are stable, with their basins of attraction separated by an unstable interior equilibrium 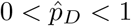 whose value is given by the shade of brown. Each panel is based on 100×100 parameter combinations. Other fixed parameters: *e* = 1, *m* = 0, other fitness costs equal to zero.

**Figure S5:**
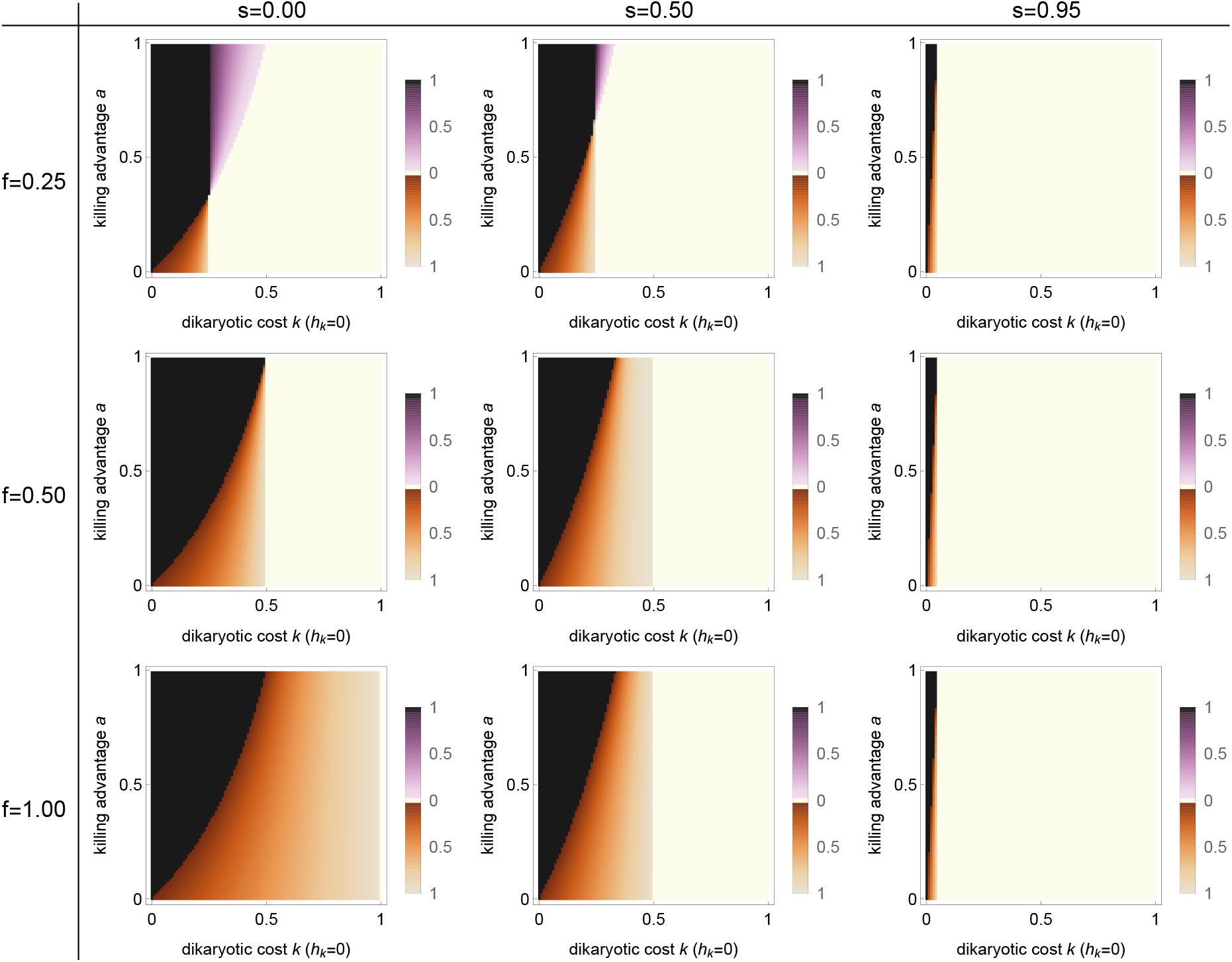
Bifurcation analysis of the *Podospora* model with recessive (*h*_*k*_ = 0) dikaryotic fitness costs *k*. The bifurcation parameters are dikaryotic fitness costs *k*, killing advantage *a*, selfing rate *s* and probability of first-division segregation *f*. Extinction 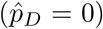 and fixation 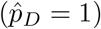 of the killer allele *D* always constitute equilibria. Additionally, one interior equilibrium is possible. Parameter regions are color coded as follows: **white**, *D* cannot invade and 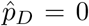 is a globally stable equilibrium; **black**, *D* can invade and reaches fixation, 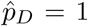 is a globally stable equilibrium; **purple**, *D* can invade but cannot reach fixation and coexists with the non-killing allele *d* at a globally stable interior equilibrium 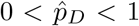, whose value is given by the shade of purple; **brown**, the two boundary equilibria 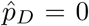 and 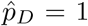 are stable, with their basins of attraction separated by an unstable interior equilibrium 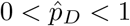 whose value is given by the shade of brown. Each panel is based on 100×100 parameter combinations. Other fixed parameters: *e* = 1, *m* = 0, other fitness costs equal to zero.

**Figure S6:**
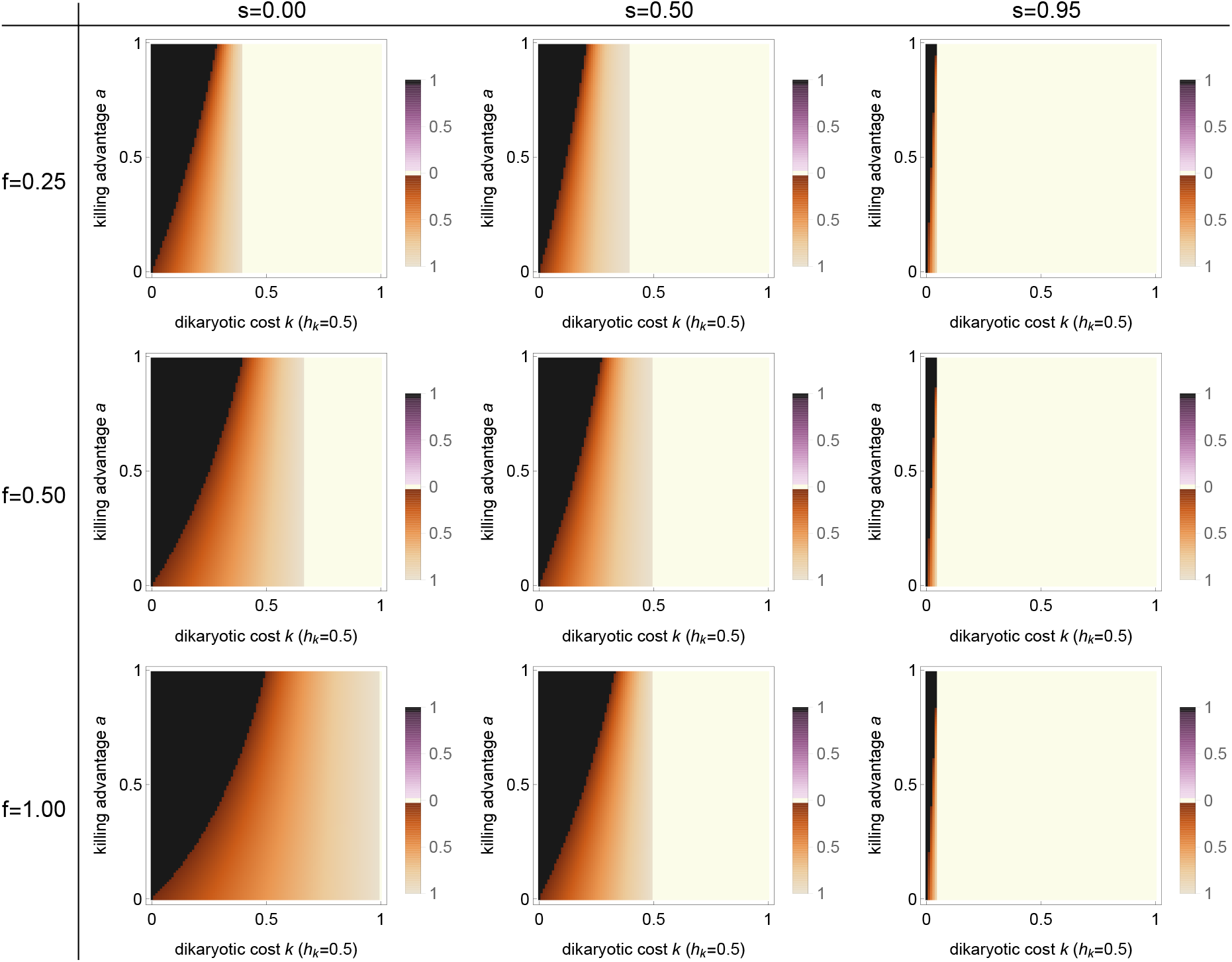
Bifurcation analysis of the *Podospora* model with additive (*h*_*k*_ = 0.5) dikaryotic fitness costs *k*. The bifurcation parameters are dikaryotic fitness costs *k*, killing advantage *a*, selfing rate *s* and probability of first-division segregation *f*. Extinction 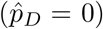 and fixation 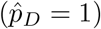 of the killer allele *D* always constitute equilibria. Additionally, one interior equilibrium is possible. Parameter regions are color coded as follows: **white**, *D* cannot invade and 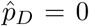 is a globally stable equilibrium; **black**, *D* can invade and reaches fixation, 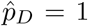 is a globally stable equilibrium; **purple**, *D* can invade but cannot reach fixation and coexists with the non-killing allele *d* at a globally stable interior equilibrium 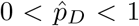, whose value is given by the shade of purple; **brown**, the two boundary equilibria 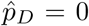 and 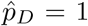 are stable, with their basins of attraction separated by an unstable interior equilibrium 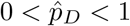 whose value is given by the shade of brown. Each panel is based on 100×100 parameter combinations. Other fixed parameters: *e* = 1, *m* = 0, other fitness costs equal to zero.

**Figure S7:**
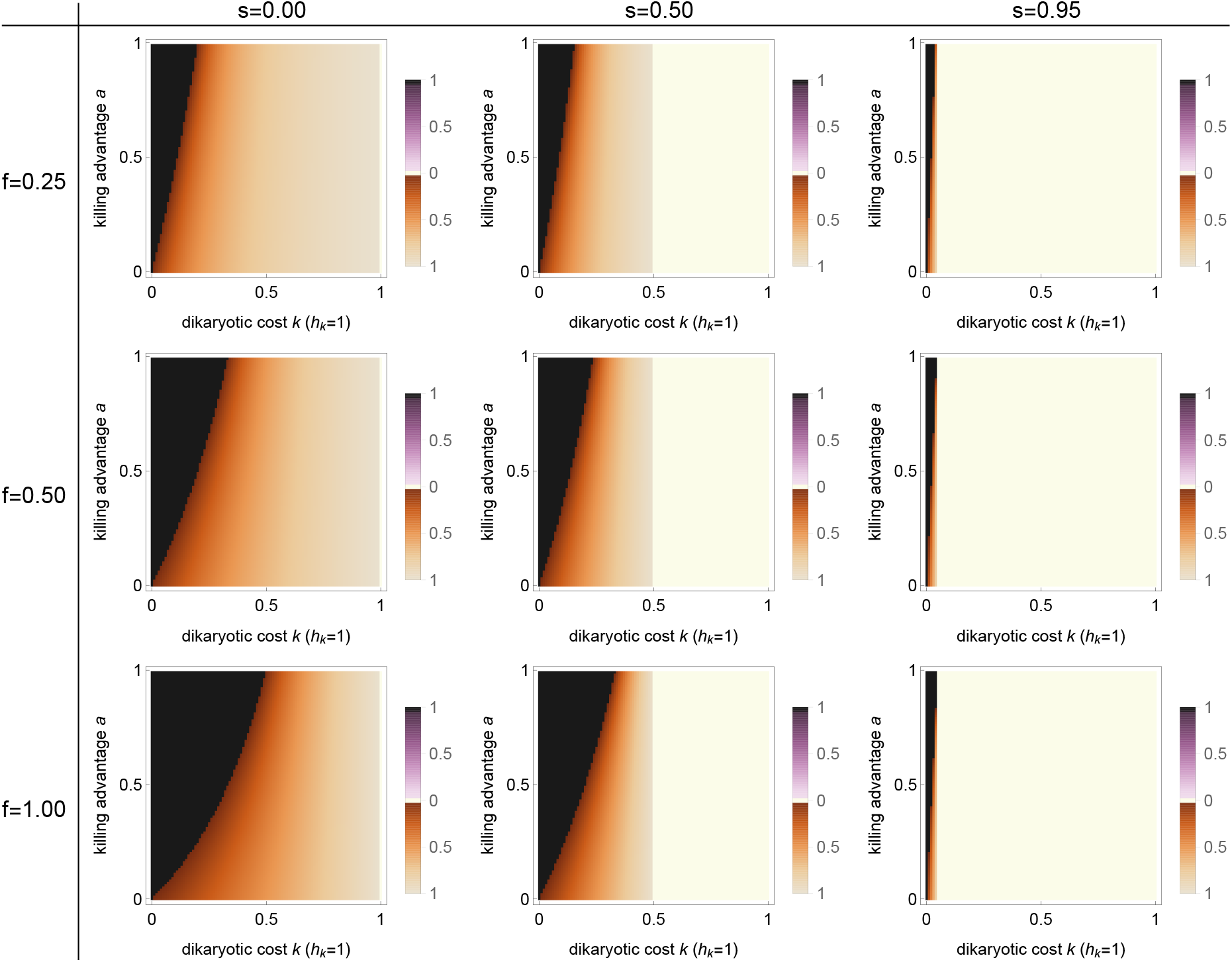
Bifurcation analysis of the *Podospora* model with dominant (*h*_*k*_ = 1) dikaryotic fitness costs *k*. The bifurcation parameters are dikaryotic fitness costs *k*, killing advantage *a*, selfing rate *s* and probability of first-division segregation *f*. Extinction 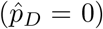 and fixation 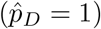 of the killer allele *D* always constitute equilibria. Additionally, one interior equilibrium is possible. Parameter regions are color coded as follows: **white**, *D* cannot invade and 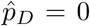 0 is a globally stable equilibrium; **black**, *D* can invade and reaches fixation, 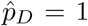 is a globally stable equilibrium; **purple**, *D* can invade but cannot reach fixation and coexists with the non-killing allele *d* at a globally stable interior equilibrium 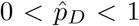, whose value is given by the shade of purple; **brown**, the two boundary equilibria 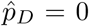 and 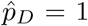 are stable, with their basins of attraction separated by an unstable interior equilibrium 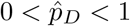 whose value is given by the shade of brown. Each panel is based on 100×100 parameter combinations. Other fixed parameters: *e* = 1, *m* = 0, other fitness costs equal to zero.

**Figure S8:**
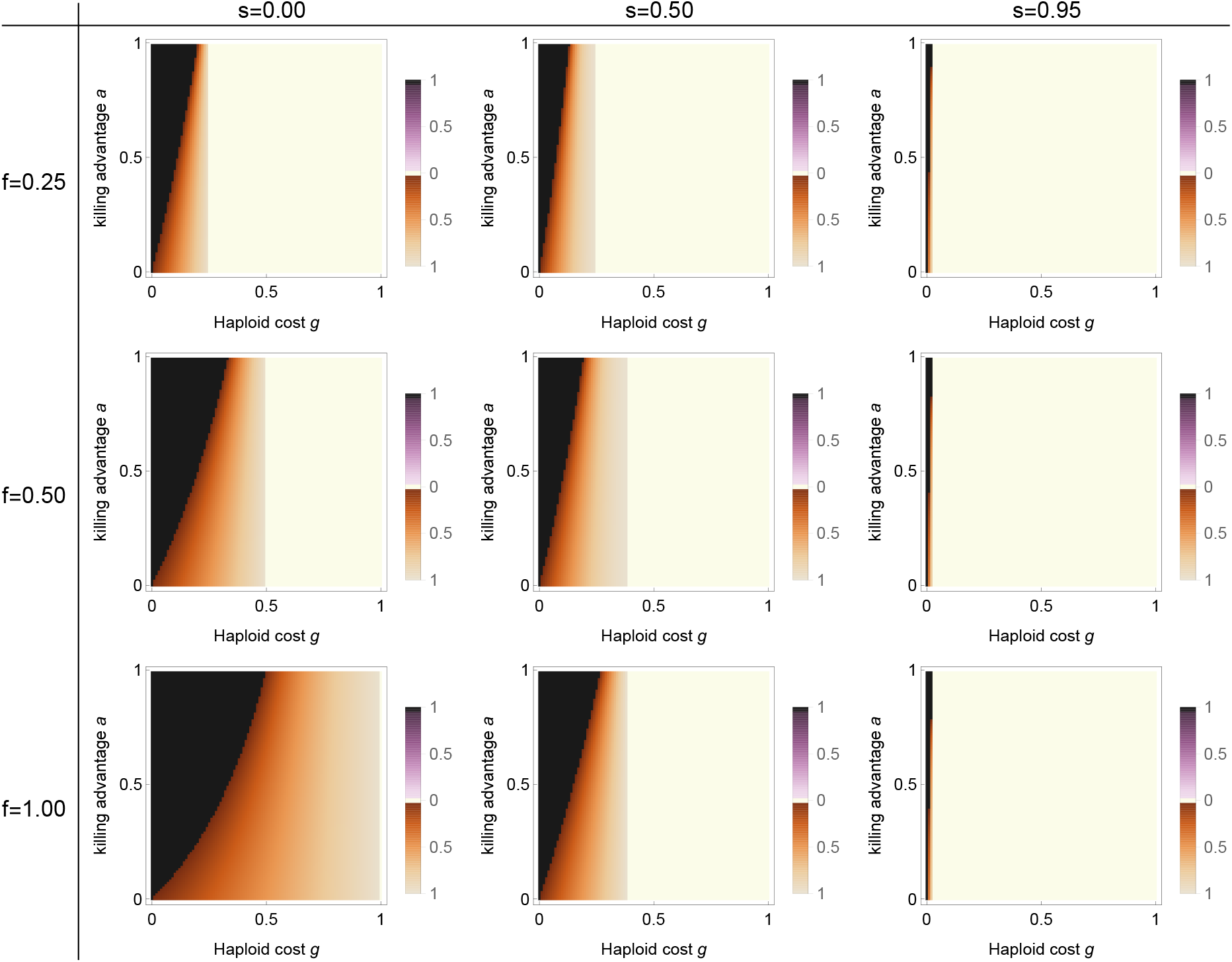
Bifurcation analysis of the *Podospora* model with haploid fitness costs *g*. The bifurcation parameters are haploid fitness costs *g*, killing advantage *a*, selfing rate *s* and probability of first-division segregation *f*. Extinction 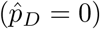 and fixation 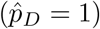 of the killer allele *D* always constitute equilibria. Additionally, one interior equilibrium is possible. Parameter regions are color coded as follows: **white**, *D* cannot invade and 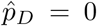 is a globally stable equilibrium; **black**, *D* can invade and reaches fixation, 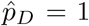 is a globally stable equilibrium; **purple**, *D* can invade but cannot reach fixation and coexists with the non-killing allele *d* at a globally stable interior equilibrium 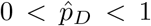, whose value is given by the shade of purple; **brown**, the two boundary equilibria 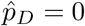 and 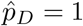 are stable, with their basins of attraction separated by an unstable interior equilibrium 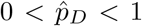 whose value is given by the shade of brown. Each panel is based on 100×100 parameter combinations. Other fixed parameters: *e* = 1, *m* = 0, other fitness costs equal to zero.

**Figure S9:**
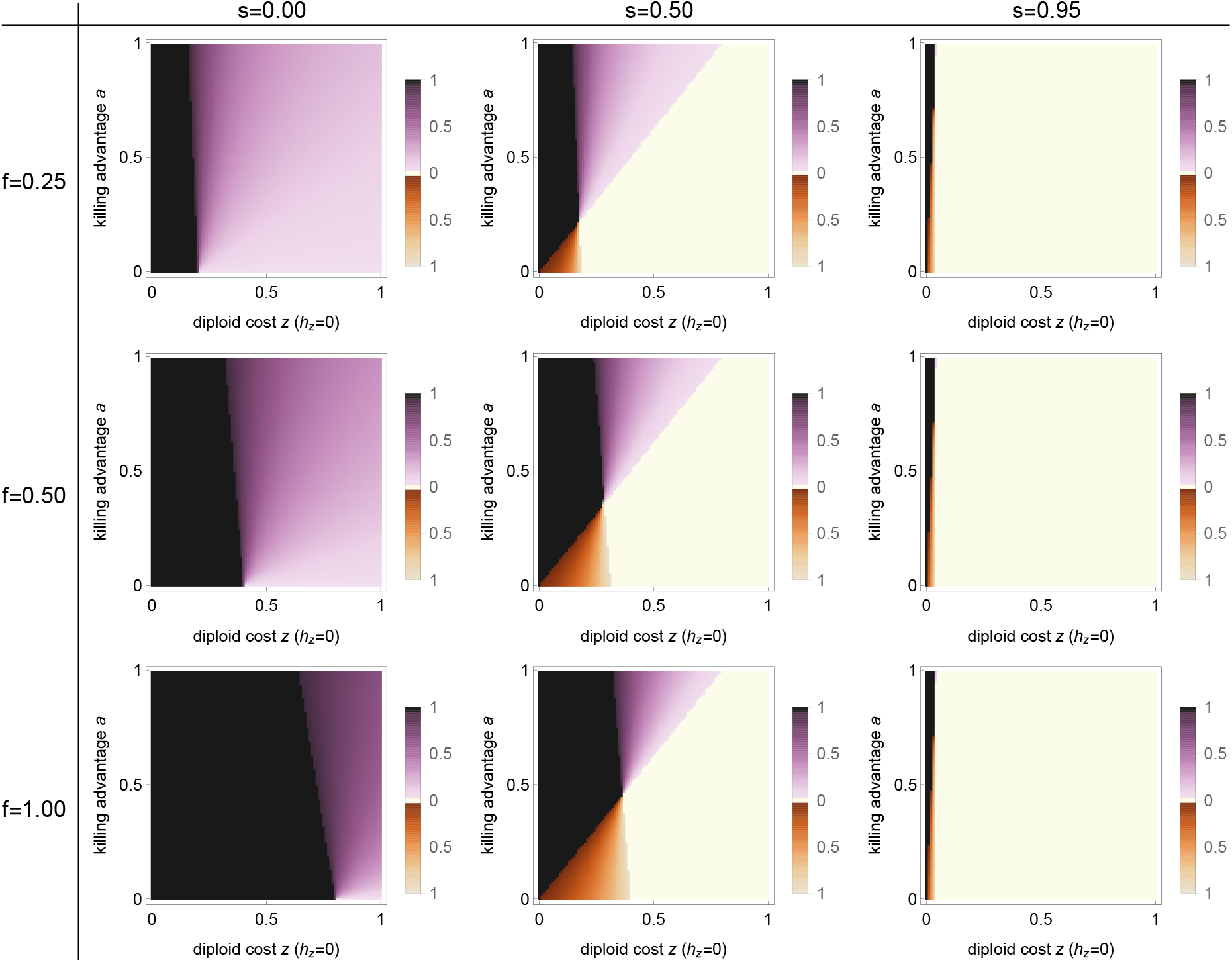
Bifurcation analysis of the *Podospora* model with incomplete killing efficiency (*e* = 0.8) and recessive (*h*_*z*_ = 0) diploid fitness costs *z*. The bifurcation parameters are diploid fitness costs *z*, killing advantage *a*, selfing rate *s* and probability of first-division segregation *f*. Extinction 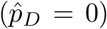 and fixation 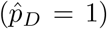 of the killer allele *D* always constitute equilibria. Additionally, one interior equilibrium is possible. Parameter regions are color coded as follows: **white**, *D* cannot invade and 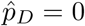 is a globally stable equilibrium; **black**, *D* can invade and reaches fixation, 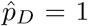 is a globally stable equilibrium; **purple**, *D* can invade but cannot reach fixation and coexists with the non-killing allele *d* at a globally stable interior equilibrium 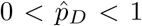, whose value is given by the shade of purple; **brown**, the two boundary equilibria 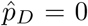 and 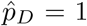 are stable, with their basins of attraction separated by an unstable interior equilibrium 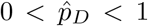 whose value is given by the shade of brown. Each panel is based on 100×100 parameter combinations. Fixed parameters: *m* = 0, other fitness costs equal to zero.

**Figure S10:**
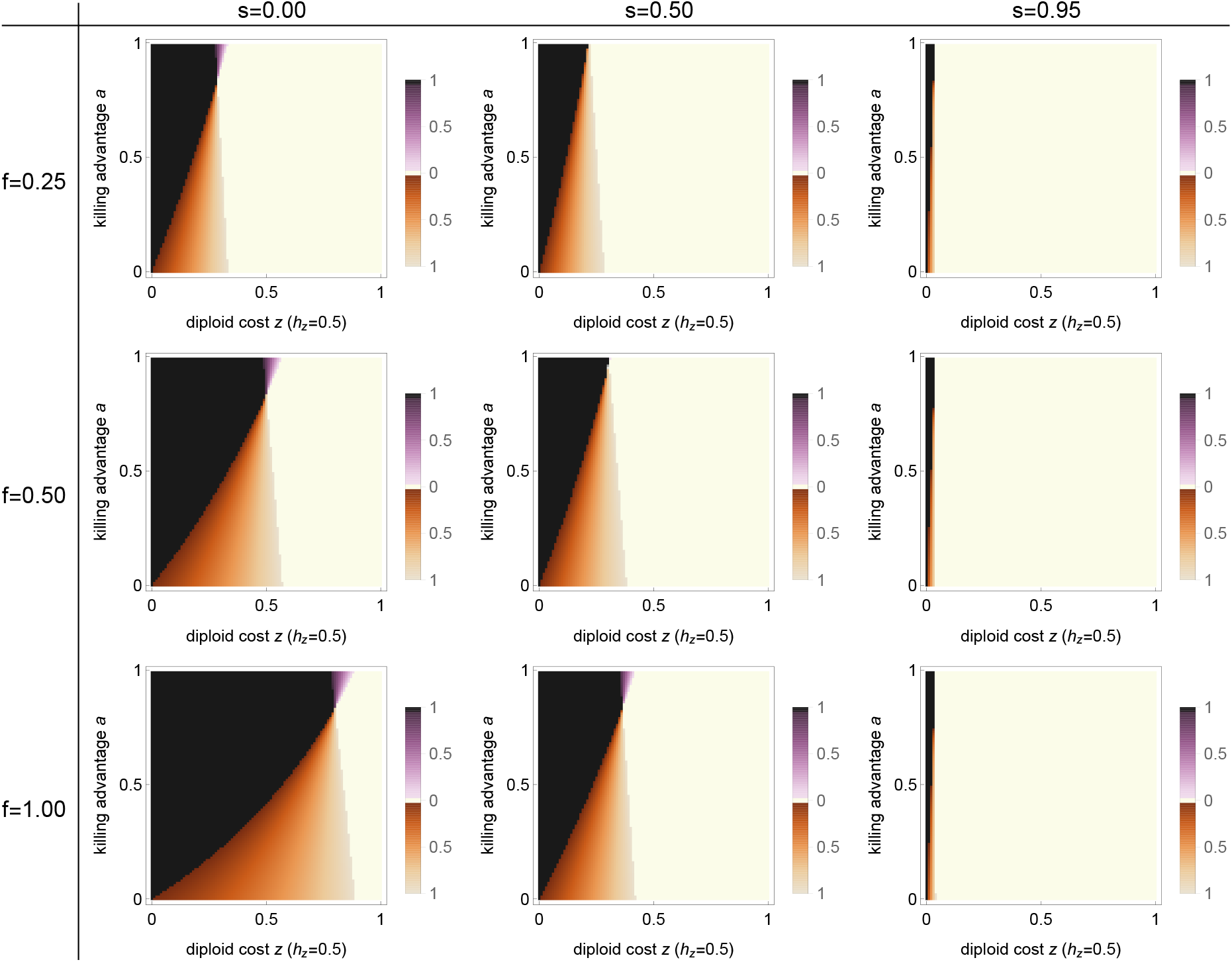
Bifurcation analysis of the *Podospora* model with incomplete killing efficiency (*e* = 0.8) and additive (*h*_*z*_ = 0.5) diploid fitness costs *z*. The bifurcation parameters are diploid fitness costs *z*, killing advantage *a*, selfing rate *s* and probability of first-division segregation *f*. Extinction 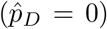 and fixation 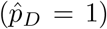 of the killer allele *D* always constitute equilibria. Additionally, one interior equilibrium is possible. Parameter regions are color coded as follows: **white**, *D* cannot invade and 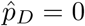 is a globally stable equilibrium; **black**, *D* can invade and reaches fixation, 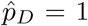 is a globally stable equilibrium; **purple**, *D* can invade but cannot reach fixation and coexists with the non-killing allele *d* at a globally stable interior equilibrium 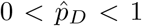, whose value is given by the shade of purple; **brown**, the two boundary equilibria 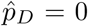 and 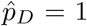 are stable, with their basins of attraction separated by an unstable interior equilibrium 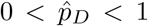 whose value is given by the shade of brown. Each panel is based on 100×100 parameter combinations. Fixed parameters: *m* = 0, other fitness costs equal to zero.

**Figure S11:**
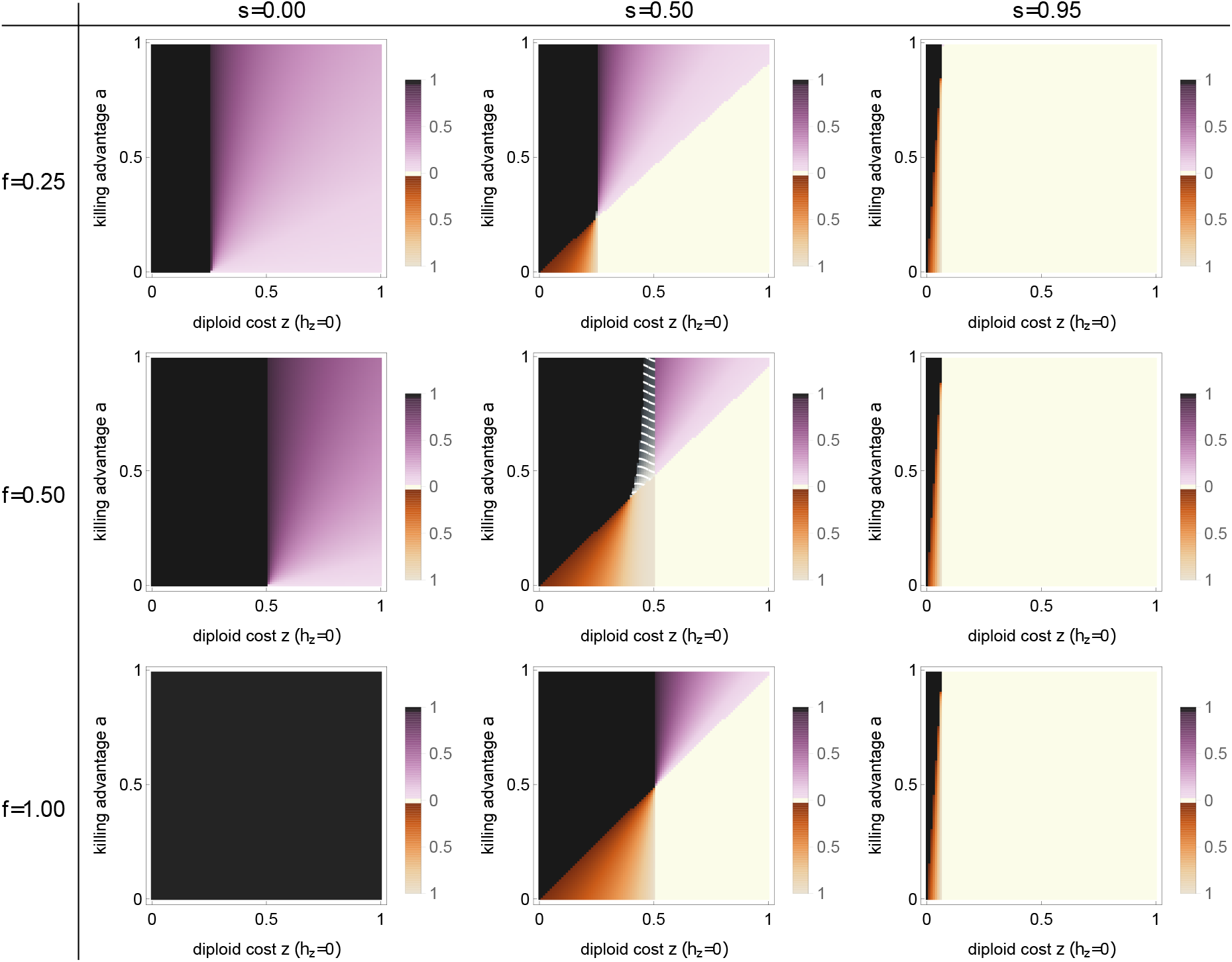
Bifurcation analysis of the *Podospora* model with monokaryons in 5% of asci (*m* = 0.05) and recessive (*h*_*z*_ = 0) diploid fitness costs *z*. The bifurcation parameters are diploid fitness costs *z*, killing advantage *a*, selfing rate *s* and probability of first-division segregation *f*. Extinction 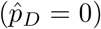 and fixation 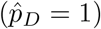 of the killer allele *D* always constitute equilibria. Additionally, one interior equilibria is possible. Parameter regions are color coded as follows: **white**, *D* cannot invade and 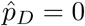 is a globally stable equilibrium; **black**, *D* can invade and reaches fixation, 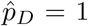 is a globally stable equilibrium; **purple**, *D* can invade but cannot reach fixation and coexists with the non-killing allele *d* at a globally stable interior equilibrium 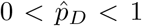, whose value is given by the shade of purple; **brown**, the two boundary equilibria 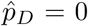 and 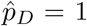 are stable, with their basins of attraction separated by an unstable interior equilibrium 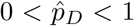 whose value is given by the shade of brown; **gray with white stripes**, two interior equilibria exist, the equilibrium with the lower value is stable, meaning that *D* can invade and coexist with *d* at a stable interior equilibrium, whose value is given by the shade of gray. Each panel is based on 100×100 parameter combinations. Fixed parameters: *e* = 1, other fitness costs equal to zero.

**Figure S12:**
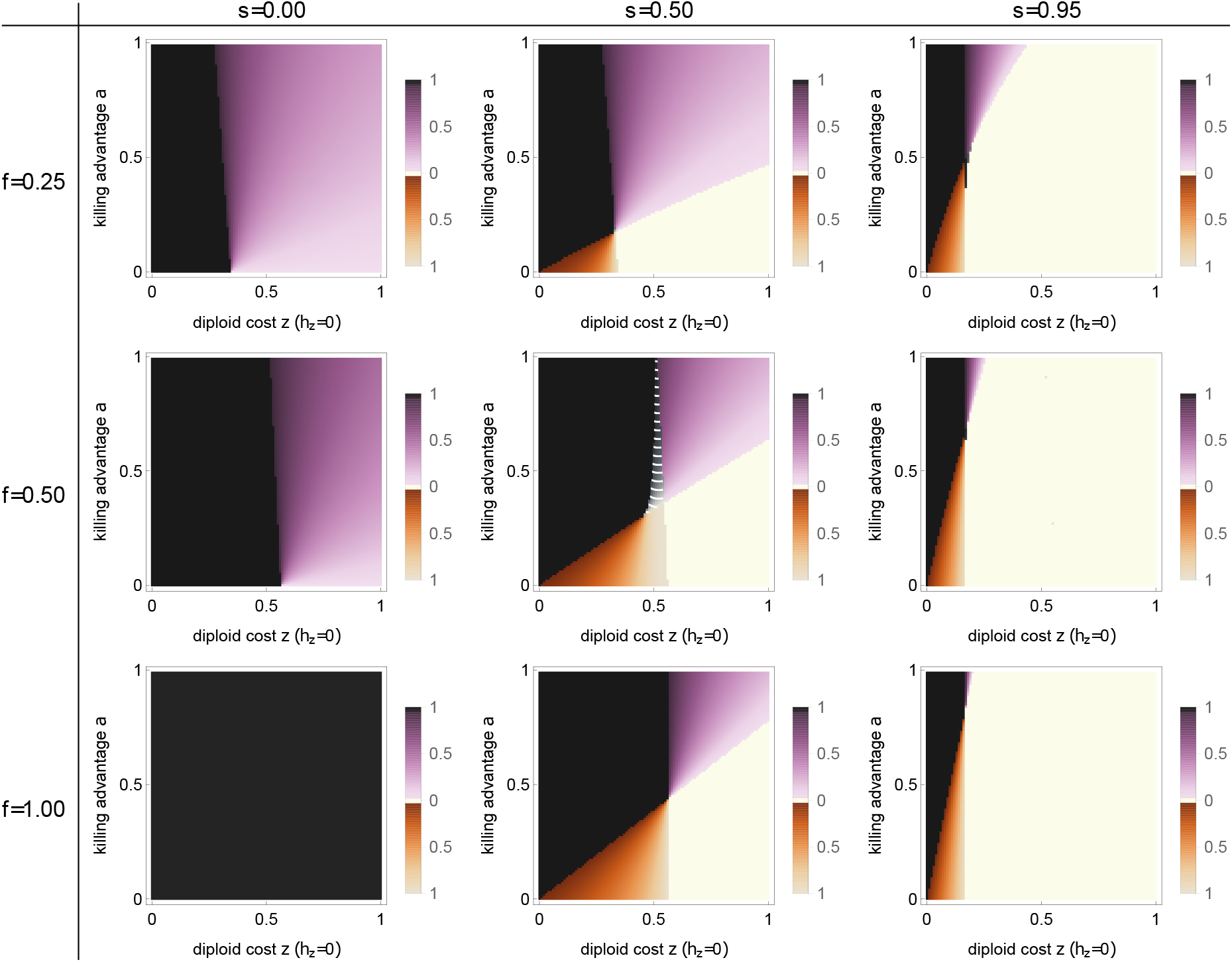
Bifurcation analysis of the *Podospora* model with monokaryons in 50% of asci (*m* = 0.5) and recessive (*h*_*z*_ = 0) diploid fitness costs *z*. The bifurcation parameters are diploid fitness costs *z*, killing advantage *a*, selfing rate *s* and probability of first-division segregation *f*. Extinction 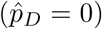 and fixation 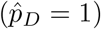 of the killer allele *D* always constitute equilibria. Additionally, one interior equilibrium is possible. Parameter regions are color coded as follows: **white**, *D* cannot invade and 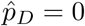 is a globally stable equilibrium; **black**, *D* can invade and reaches fixation, 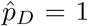 is a globally stable equilibrium; **purple**, *D* can invade but cannot reach fixation and coexists with the non-killing allele *d* at a globally stable interior equilibrium 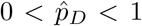, whose value is given by the shade of purple; **brown**, the two boundary equilibria 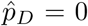 and 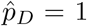 are stable, with their basins of attraction separated by an unstable interior equilibrium 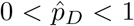 whose value is given by the shade of brown; **gray with white stripes**, two interior equilibria exist, the equilibrium with the lower value is stable, meaning that *D* can invade and coexist with *d* at a stable interior equilibrium, whose value is given by the shade of gray. Each panel is based on 100×100 parameter combinations. Fixed parameters: *e* = 1, other fitness costs equal to zero.

**Figure S13:**
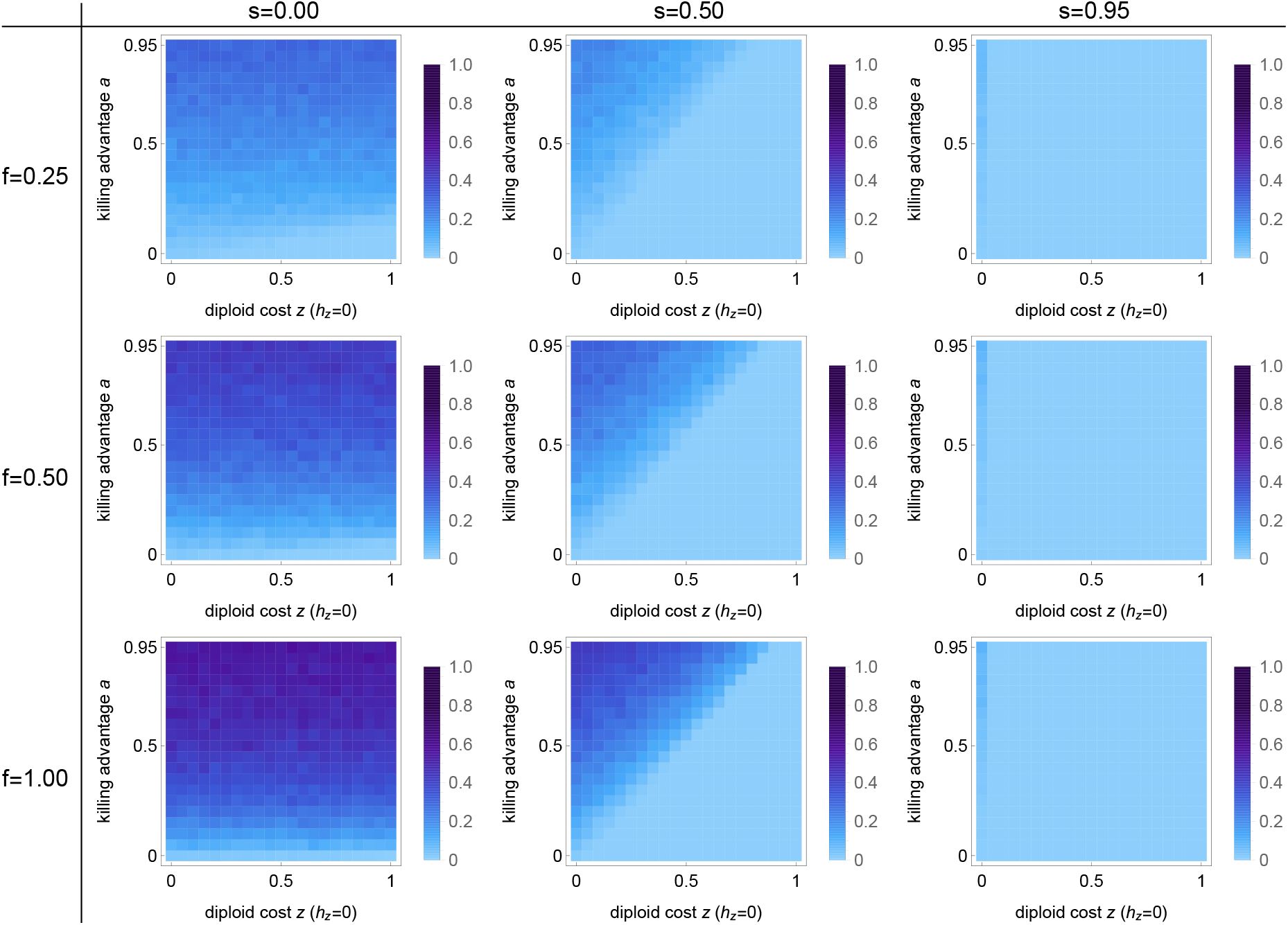
Invasion probability of a spore-killing allele *D* for the *Podospora* model with recessive (*h*_*z*_ = 0) diploid fitness costs *z*. Parameters are the fitness costs *z*, the killing advantage *a*, the selfing rate *s*, and the probability of first-division segregation *f*. Each panel consists of 21×21 parameter combinations and shades of blue indicates the invasion probability estimated from 10^3^ stochastic Wright-Fisher simulation runs with a population size of 1000. Other parameters as in Figure 5.

**Figure S14:**
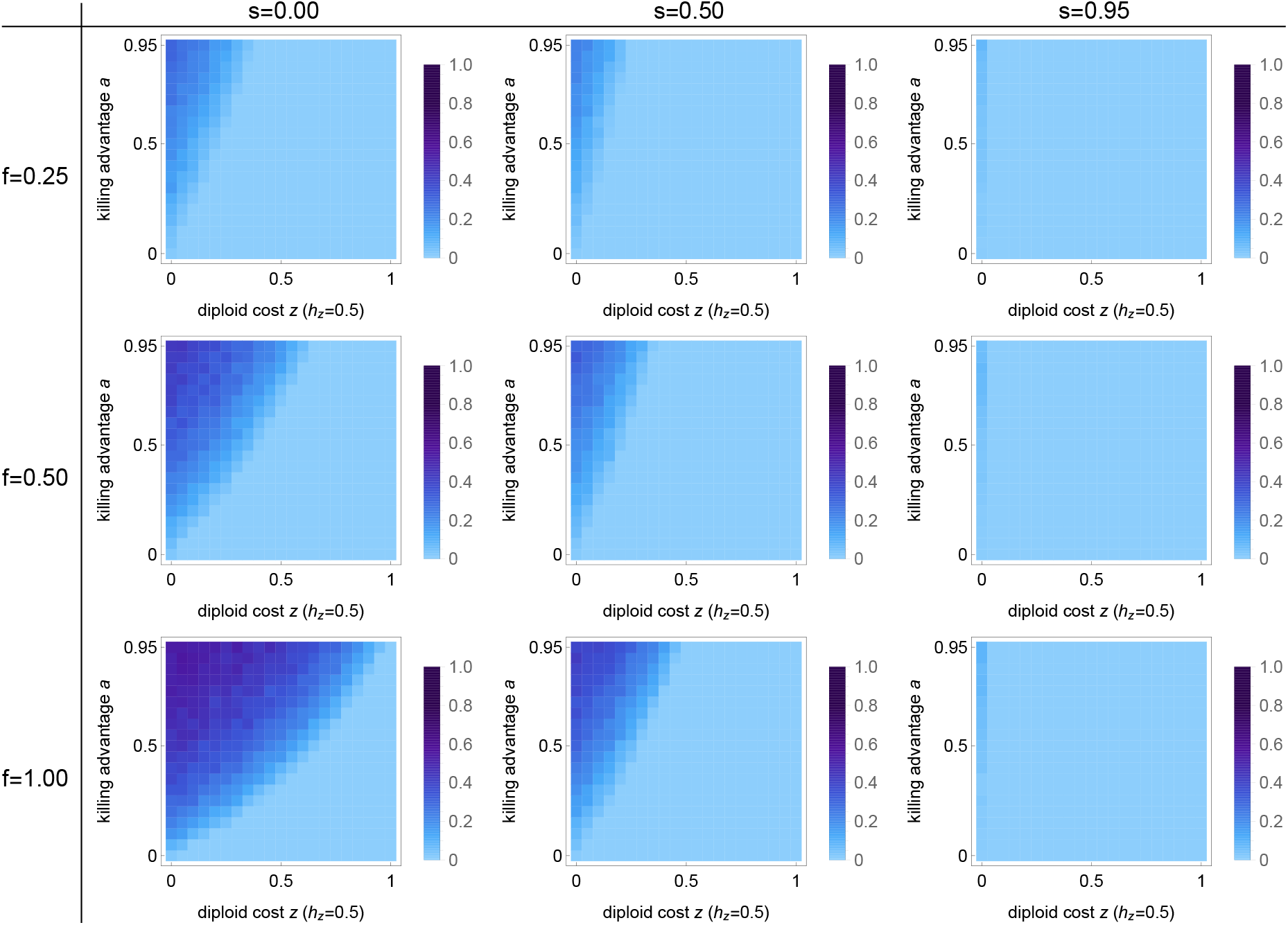
Invasion probability of a spore-killing allele *D* for the *Podospora* model with additive (*h*_*z*_ = 0.5) diploid fitness costs *z*. Parameters are the fitness costs *z*, the killing advantage *a*, the selfing rate *s* and the rate of first-division segregation *f*. Each panel consists of 21×21 parameter combinations and shades of blue indicates the invasion probability estimated from 10^3^ stochastic Wright-Fisher simulation runs with a population size of 1000. Other parameters as in Figure S3.

**Figure S15:**
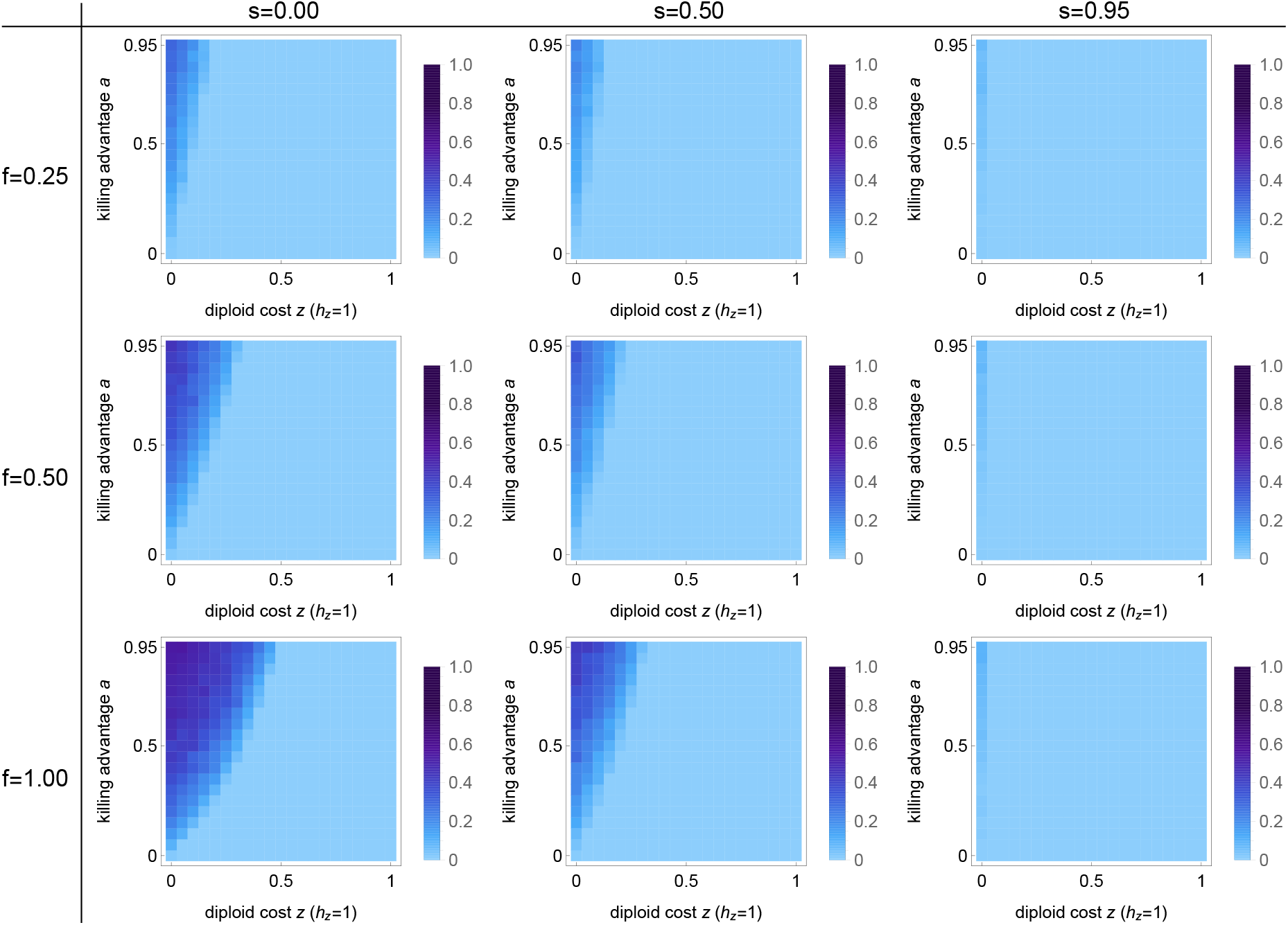
Invasion probability of a spore-killing allele *D* for the *Podospora* model with dominant (*h*_*z*_ = 1) diploid fitness costs *z*. Parameters are the fitness costs *z*, the killing advantage *a*, the selfing rate *s* and the rate of first-division segregation *f*. Each panel consists of 21×21 parameter combinations and shades of blue indicate the invasion probability estimated from 10^3^ stochastic Wright-Fisher simulation runs with a population size of 1000. Other parameters as in Figure S4.

**Figure S16:**
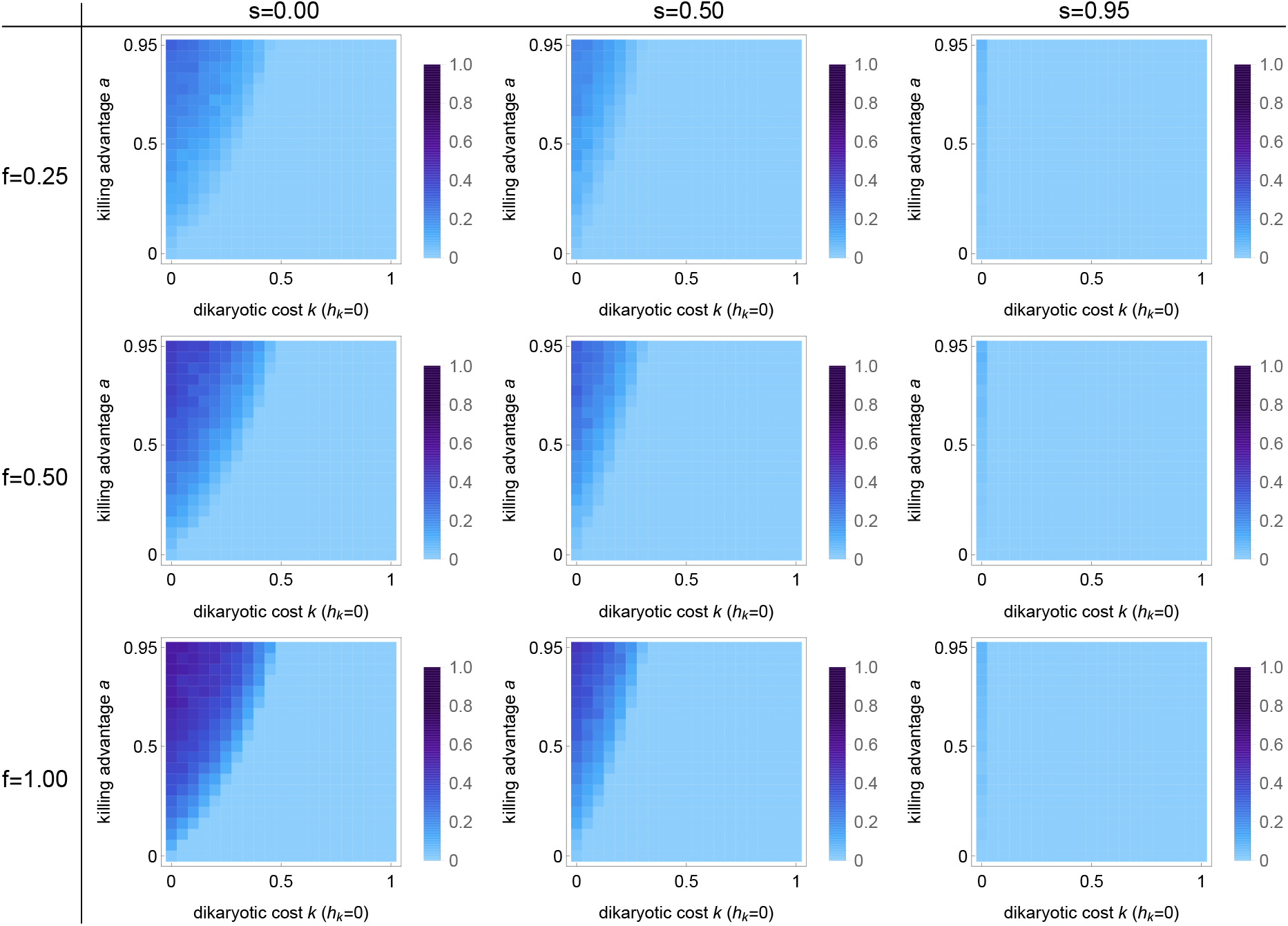
Invasion probability of a spore-killing allele *D* for the *Podospora* model with recessive (*h*_*k*_ = 0) dikaryotic fitness costs *k*. Parameters are the fitness costs *k*, the killing advantage *a*, the selfing rate *s* and the rate of first-division segregation *f*. Each panel consists of 21×21 parameter combinations and shades of blue indicate the invasion probability estimated from 10^3^ stochastic Wright-Fisher simulation runs with a population size of 1000. Other parameters as in Figure S5.

**Figure S17:**
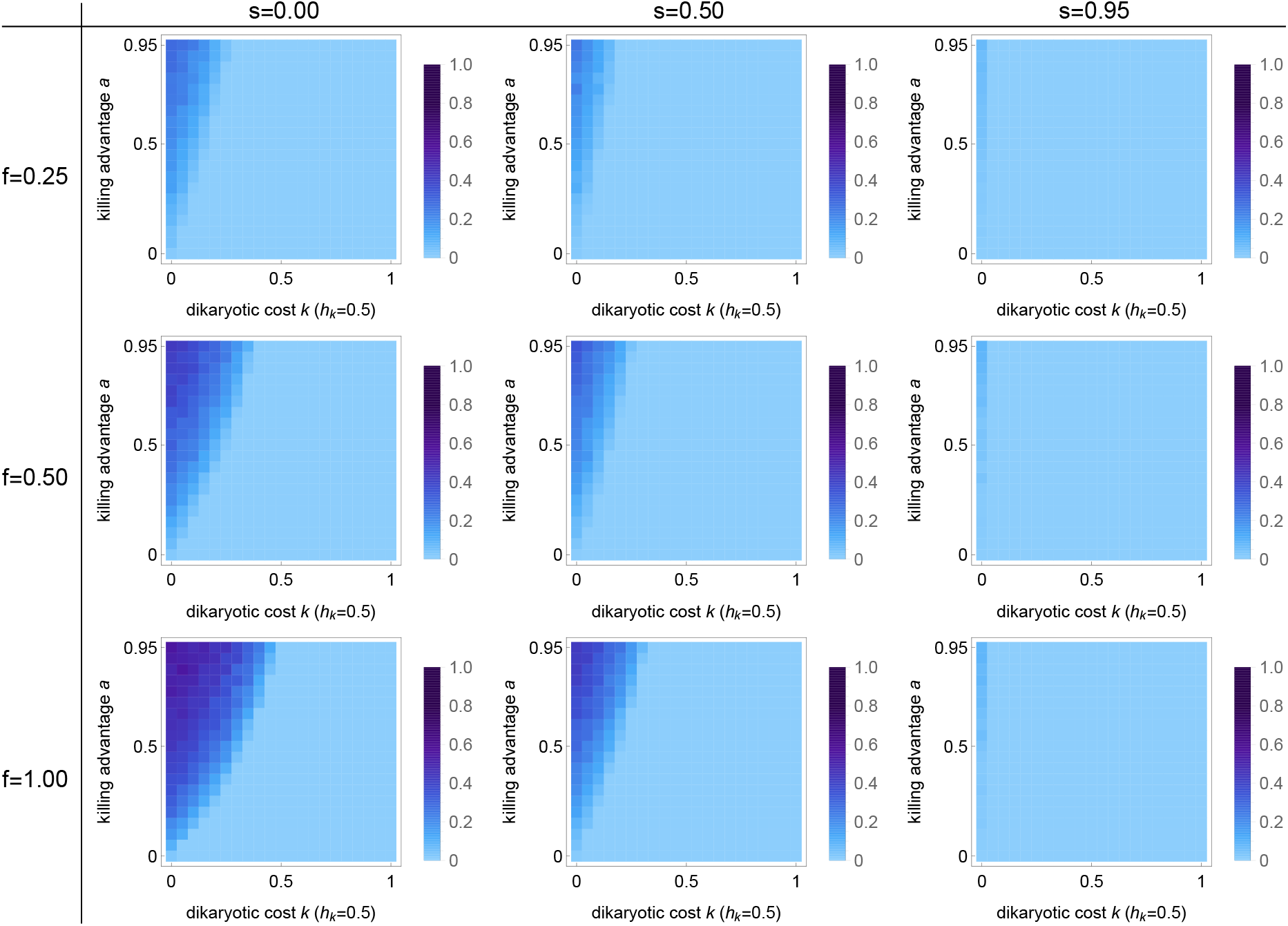
Invasion probability of a spore-killing allele *D* for the *Podospora* model with additive (*h*_*k*_ = 1*/*2) dikaryotic fitness costs *k*. Parameters are the fitness costs *k*, the killing advantage *a*, the selfing rate *s* and the rate of first-division segregation *f*. Each panel consists of 21×21 parameter combinations and shades of blue indicate the invasion probability estimated from 10^3^ stochastic Wright-Fisher simulation runs with a population size of 1000. Other parameters as in Figure S6.

**Figure S18:**
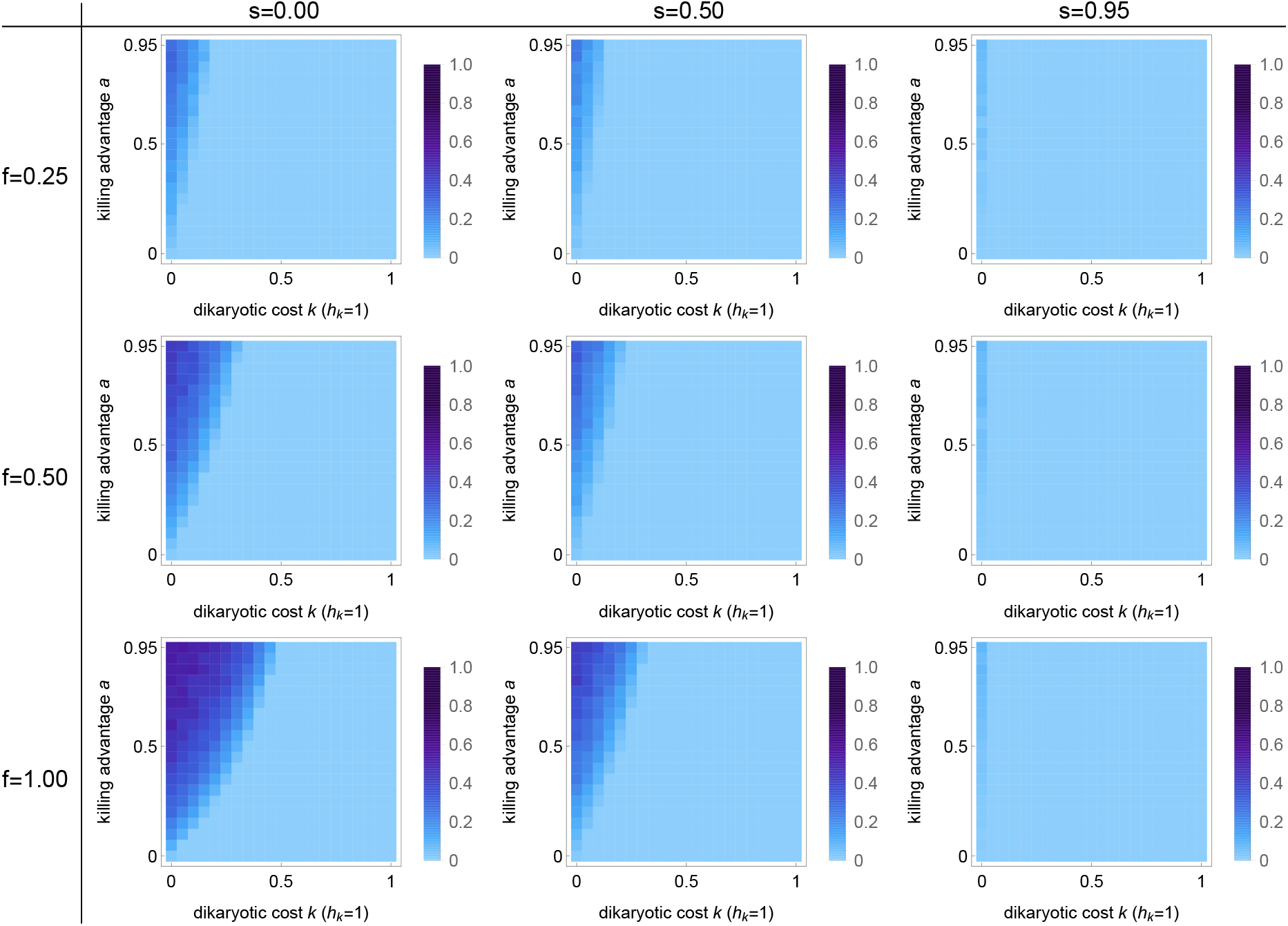
Invasion probability of a spore-killing allele *D* for the *Podospora* model with dominant (*h*_*k*_ = 1) dikaryotic fitness costs *k*. Parameters are the fitness costs *k*, the killing advantage *a*, the selfing rate *s* and the rate of first-division segregation *f*. Each panel consists of 21×21 parameter combinations and shades of blue indicate the invasion probability estimated from 10^3^ stochastic Wright-Fisher simulation runs with a population size of 1000. Other parameters as in Figure S7.

**Figure S19:**
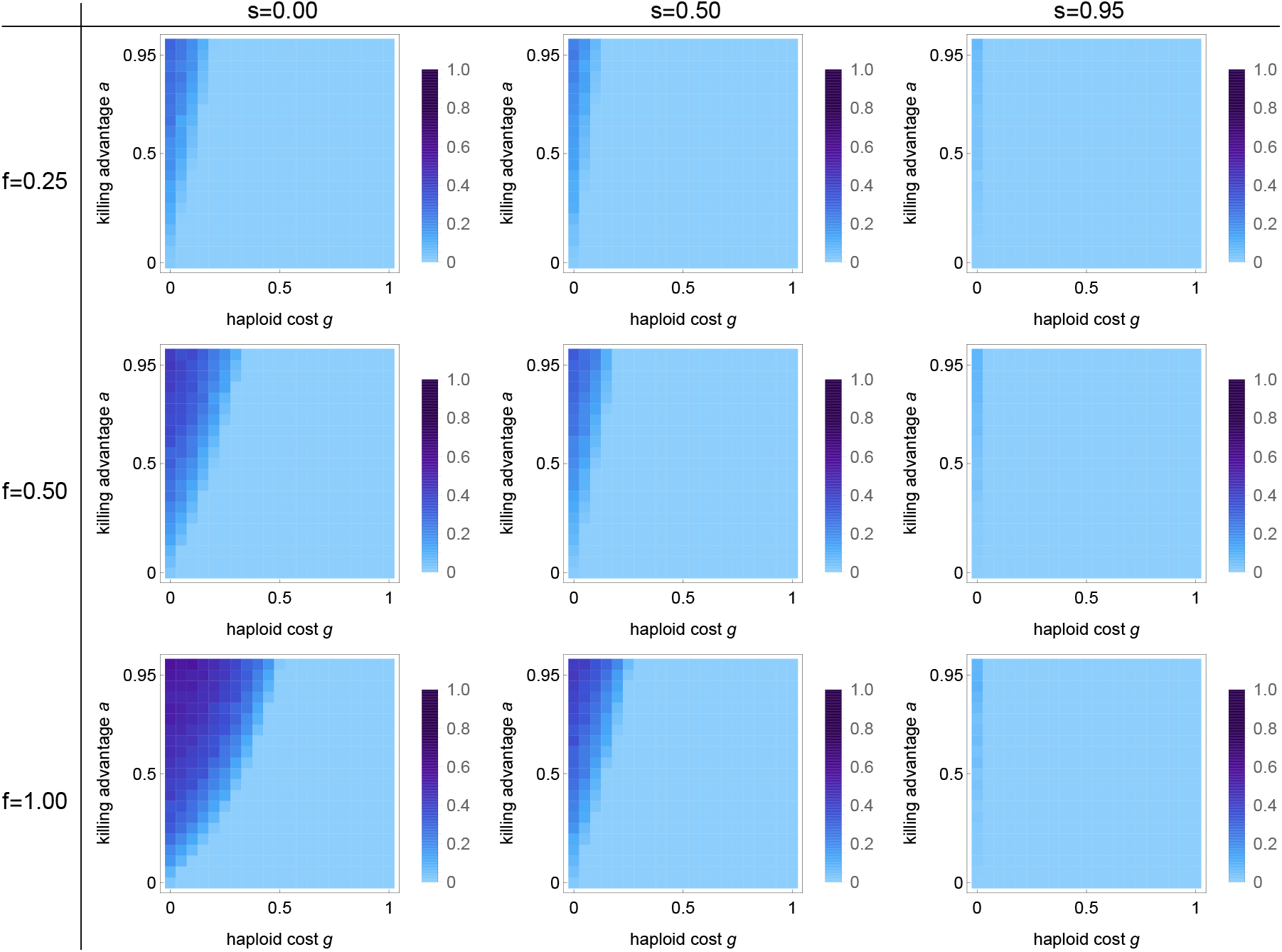
Invasion probability of a spore-killing allele *D* for the *Podospora* model with haploid fitness costs *g*. Parameters are the fitness costs *g*, the killing advantage *a*, the selfing rate *s* and the rate of first-division segregation *f*. Each panel consists of 21×21 parameter combinations and shades of blue indicate the invasion probability estimated from 10^3^ stochastic Wright-Fisher simulation runs with a population size of 1000. Other parameters as in Figure S8.

**Figure S20:**
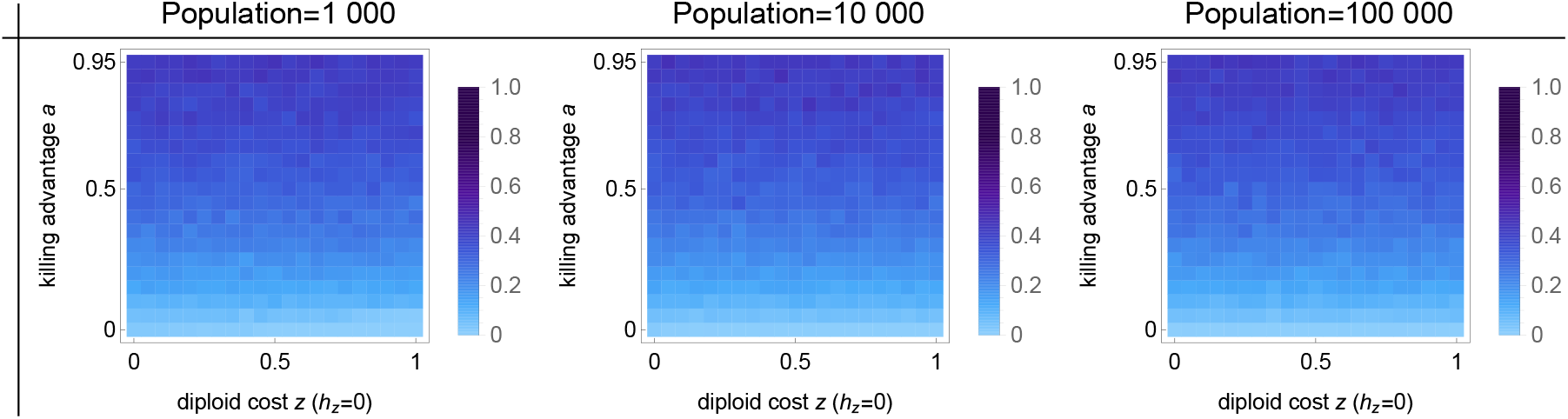
Effect of population size on invasion probability of a spore-killing allele *D* with recessive diploid costs *z*. Parameters are the recessive (*h*_*z*_ = 0) fitness costs *z*, the killing advantage *a*, and population size.The selfing rate *s* is fixed to zero and the rate of first-division segregation *f* to 0.50. Each panel consists of 21×21 parameter combinations and shades of blue indicate the invasion probability estimated from 10^3^ stochastic Wright-Fisher simulation runs.

**Figure S21:**
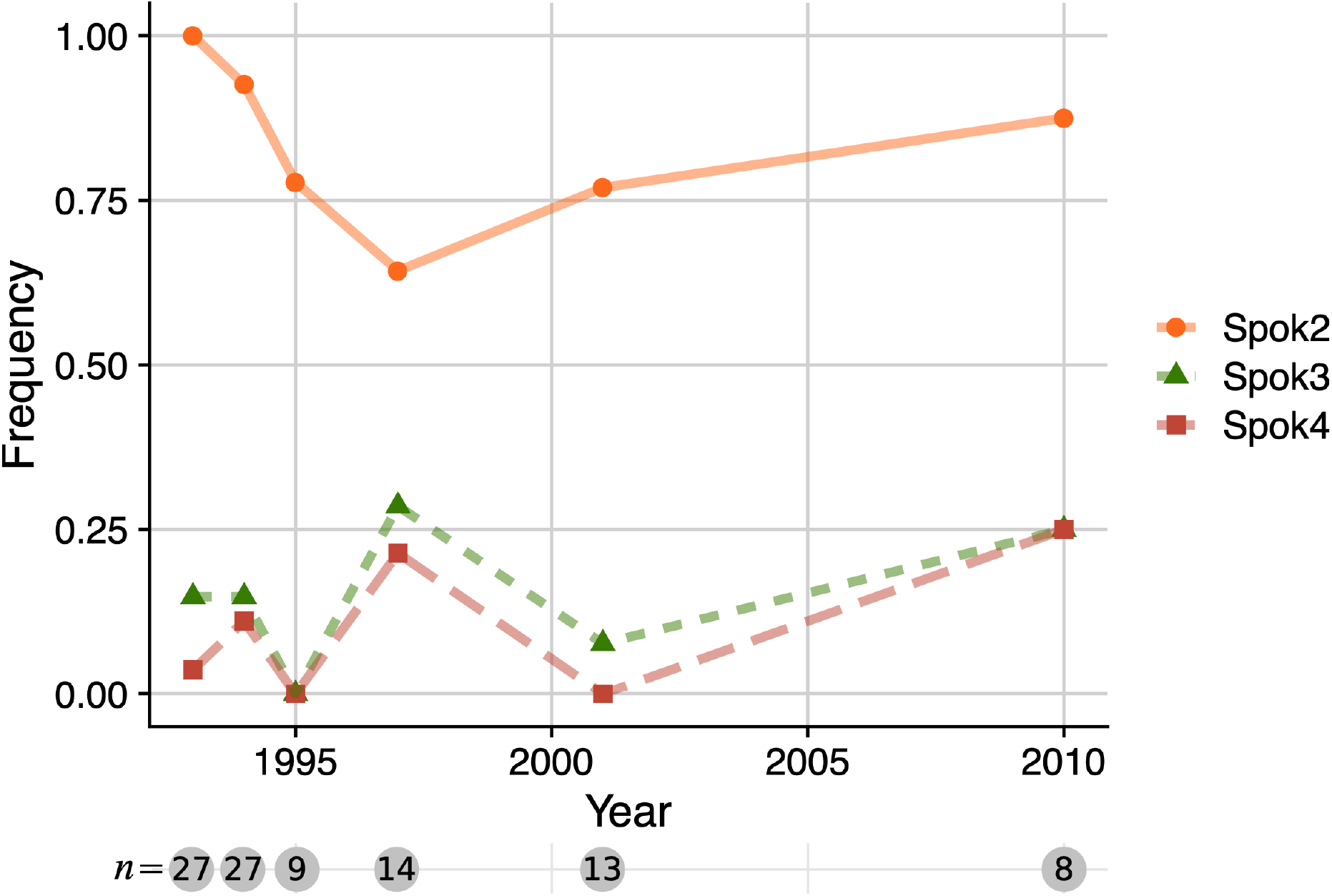
Frequency of *Spok2, Spok3* and *Spok4* in the Wageningen population of *Podospora anserina*. Individuals of *P. anserina* have been collected around Wageningen, The Netherlands, at irregular intervals between 1993 and 2010 (van der Gaag, 2005). The genome of each individual might contain any number and combination of *Spok* genes, from no *Spok* gene to all three of them. The *Spok3* and *Spok4* genes are embedded within a larger haplotype called the ‘*Spok* block’. Thus, while *Spok3* and *Spok4* can occur on their own, their occurrence seems to be tightly linked. We used published and unpublished genomic data to determine the *Spok* gene content of each individual, and verified that spore killing phenotypes followed expected patterns (Vogan et al., 2019). The number *n* gives the sample size in each year.

**Figure S22:**
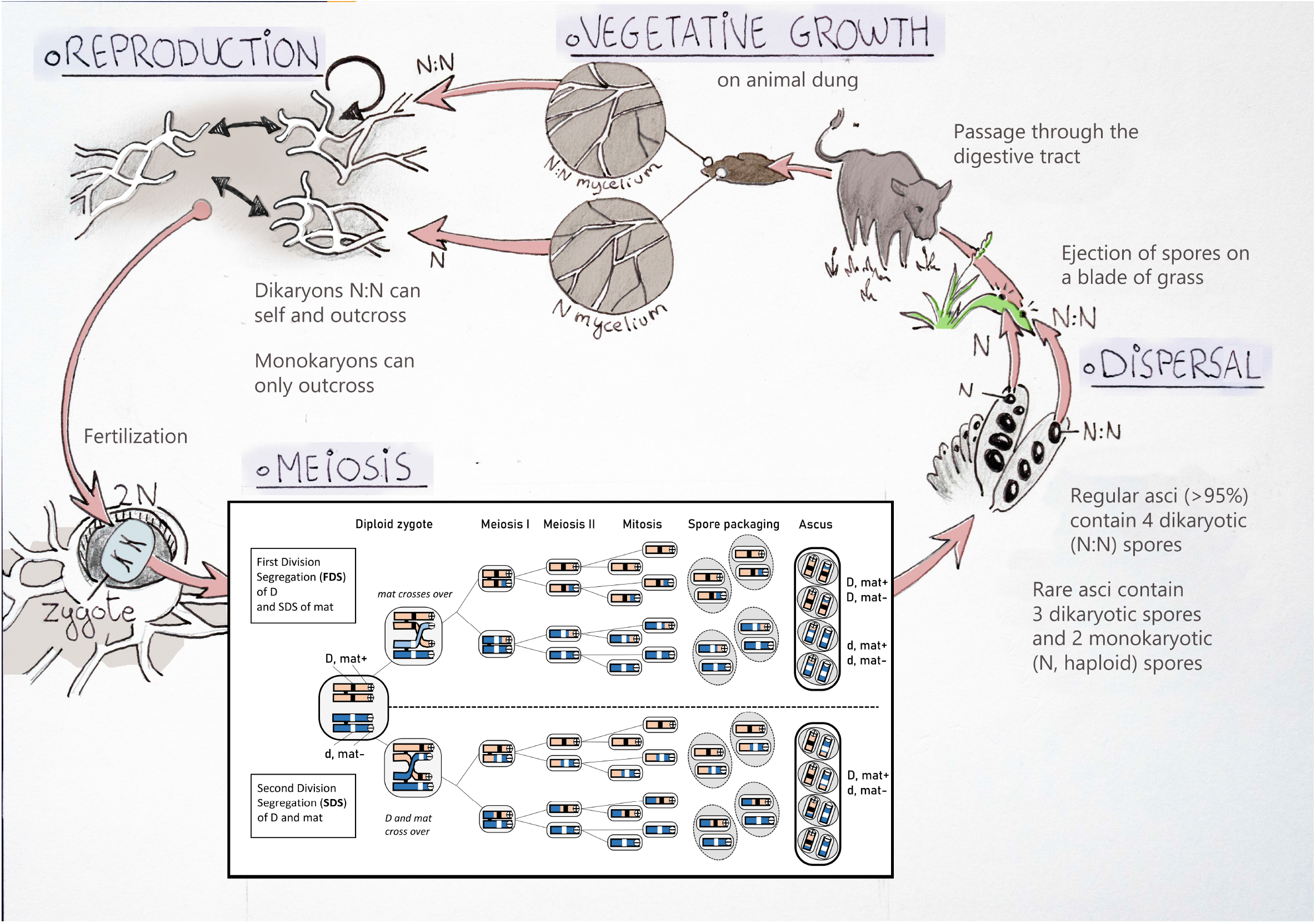
Simplified life cycle of *Podospora anserina* and meiosis diagram. The meiosis diagram is shown for the spore killer *D* and its sensitive counterpart *d*, as well as for the mating type locus with alleles *mat*+ and *mat* −. For simplicity, the spore killer and the *mat*-locus are located on the same chromosome, but this does not need to be the case. The same logic applies if they are on different chromosome. In *P. anserina*, the *mat*-locus undergoes SDS in almost all meiosis events, allowing self-fertility of the dikaryotic spores. The spore killer undergoes FDS or SDS with various probabilities depending on the position of the gene on the chromosome.

## Appendix 1: Invasion probability of a simple spore killer

### Purpose

In this appendix, we give the details for the model presented in Section 3.3. The calculations are adapted from Desai and Fisher (2007), and give an approximation for the invasion probability of a spore killer with 100% killing efficiency but no killing advantage in a randomly-mating population (i.e., no selfing), starting from a single copy. The spore killer is said to have ‘invaded’ when it has reached a sufficient copy number that its dynamics is determined predominantly by the deterministic selective advantage it obtains from killing, rather than by random drift.

### Heuristic calculation

As before, let *p*_*D*_ denote the frequency of the spore killer, which we assume to be small (*p*_*D*_ ≈ 0). Setting *a* = 0, *z* = 0, *g* = 0 and *e* = 1 in Eq. (1), we find that the selective advantage of the spore killer is

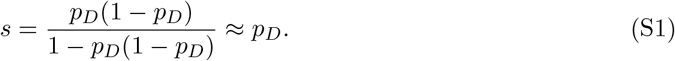

Let *N* denote the total population size and *n* the absolute number of copies of the spore killer so that *p*_*D*_ = *n/N*. We now consider a typical change in the allele’s copy number due to drift over the succeeding *n* generations, and compare it to the expected change in copy number due to positive selection on the allele. The reason we consider *n* generations is that, starting from *n* copies of the allele, the standard deviation of the fluctuation in allele copy number across *n* generations that is due to random drift is ∼ *n* (Desai and Fisher, 2007). Therefore, *n* generations is the timescale for possible extinction of the allele due to random drift. Counteracting this possibility of random loss of the spore killer allele is deterministic selection in favor of it. Across the same *n* generations, the spore killer has an expected increase in copy number due to positive selection of approximately *ns* = *np*_*D*_ = *n*^2^*/N* copies (indeed, slightly more because the selective advantage comes to exceed *s* = *p*_*D*_ as the spore killer increases in copy number), for a total expected gain of *n*^3^*/N* copies. Therefore, the deterministic force pushing the spore killer up in copy number dominates the random force that could push it down in copy number when *n*^3^*/N* > *n*, i.e., when 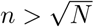. Thus, the spore killer can be said to have invaded when it has attained 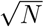 copies. Because its dynamics before this point is dominated by drift, the probability that it attains 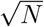 copies having started as a single copy is approximately 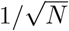 (Crow and Kimura, 1970). As shown in Fig. 6a, this approximation accords well with estimates obtained from simulations of the model.

### Comparison with a recessive beneficial allele

Consider a recessive beneficial allele *D* with selection coefficient 1 (i.e., the relative fitnesses of the *dd, Dd*, and *DD* genotypes are 1, 1, and 2, respectively; the relevant comparison is to the case of a spore killer with killing efficiency *e* = 1, as considered in the subsection above). From an initial frequency of *p*_*D*_, the change in frequency of the recessive beneficial mutation across one generation is

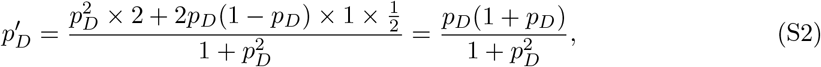

so that the allele’s selective advantage

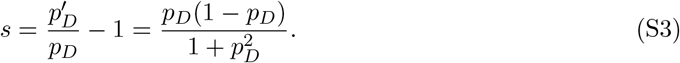

When *p*_*D*_ ≪ 1 (the relevant case for invasion), then

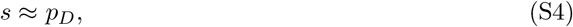

as for the spore killer considered above Eq. (S1). Thus, although they are not identical, the selective advantages of the spore killer and the recessive beneficial allele with selection coefficient 1 are similar when the two alleles are rare. This explains why the approximate invasion probability derived above for the spore killer resembles the fixation probability for a recessive beneficial mutation with selection coefficient 1 (Kimura, 1962, Eq. 15). It also explains why our result concerning the invasion rate of spore killers in a subdivided population, derived in Section 3.3, matches the analogous result for recessive beneficial mutations (Gale, 1990, p. 180-181).

